# New data on the gorgonopsian *Aelurognathus tigriceps* from the *Daptocephalus* Assemblage Zone (Karoo Basin, South Africa) with novel insights into the pelvic girdle, hind limbs and tail

**DOI:** 10.64898/2026.05.27.727204

**Authors:** Spencer K. Pevsner, Roger B.J. Benson, Christian F. Kammerer

## Abstract

Gorgonopsian therapsids represent a transitional condition in the evolution of synapsid locomotion and postcranial structure. However, most descriptions of gorgonopsians have focused on the skull, limiting their usefulness for informing patterns of postcranial evolution on the mammal stem. While some recent work has begun to focus on postcrania, especially the pectoral girdle and forelimbs, comparatively little data are available on the pelvic girdle, hind limbs and tail. We report a recently collected specimen of the late Permian gorgonopsian *Aelurognathus tigriceps* comprising a partial skull and well-preserved postcranial skeleton, including the near-complete series of dorsal vertebrae and ribs, complete pelvic girdle, hind limbs, feet, and a nearly complete tail. The tail includes more caudal vertebrae than any other published gorgonopsian. The material presented here provides an opportunity to better establish broader patterns of morphology in the gorgonopsian postcranial skeleton.

## Introduction

Gorgonopsians are a group of predatory non-mammalian therapsids known from the middle to late Permian (265–252 million years ago) (Kemp 2005; Kammerer & Rubidge 2022; Benoit *et al*. 2024; although see Matamales-Andreu *et al*. 2024). Their fossils are primarily known from southern Africa and Russia (Kammerer *et al*. 2023), as well as more recent discoveries from China and Spain (Liu and Yang 2022; Matamales-Andreu *et al*. 2024). Most historically-collected specimens are skulls, with comparably little known about their postcrania (Sigogneau-Russell 1989), although some recent descriptions have improved on this (Tatarinov 2004; Gebauer 2014; Sidor 2022; Bendel et al. 2023; Sidor and Mann 2024; Kulik 2025; Mann and Sidor 2025). One of the few species for which postcrania have been described is *Aelurognathus tigriceps*, whose holotype (SAM-PK-2342) comprises a skull, cervical and dorsal vertebrae, pectoral girdle, and forelimb (Broom and Haughton 1913). However, the posterior part of the skeleton remains largely unknown for this taxon. Here, we describe a new specimen of *Aelurognathus tigriceps* (SAM-PK-K10496) with extensive postcranial material, including the previously undescribed pelvis, hindlimbs, and tail.

SAM-PK-K10946 (Fig. 1) is a gorgonopsian skeleton preserved on a block of sandy siltstone. The posterior half of the specimen, containing a complete pelvic girdle, hind limbs, feet, and a nearly complete tail, is more completely preserved than the anterior half. Only the anterior two–thirds of the skull are preserved, as the posterior part of the skull and the cervical vertebrae were exposed by erosion (Roger M. H. Smith, unpublished field notes). The cervical vertebrae, forelimbs and pectoral girdle are mostly damaged or missing.

**Fig. 1.**
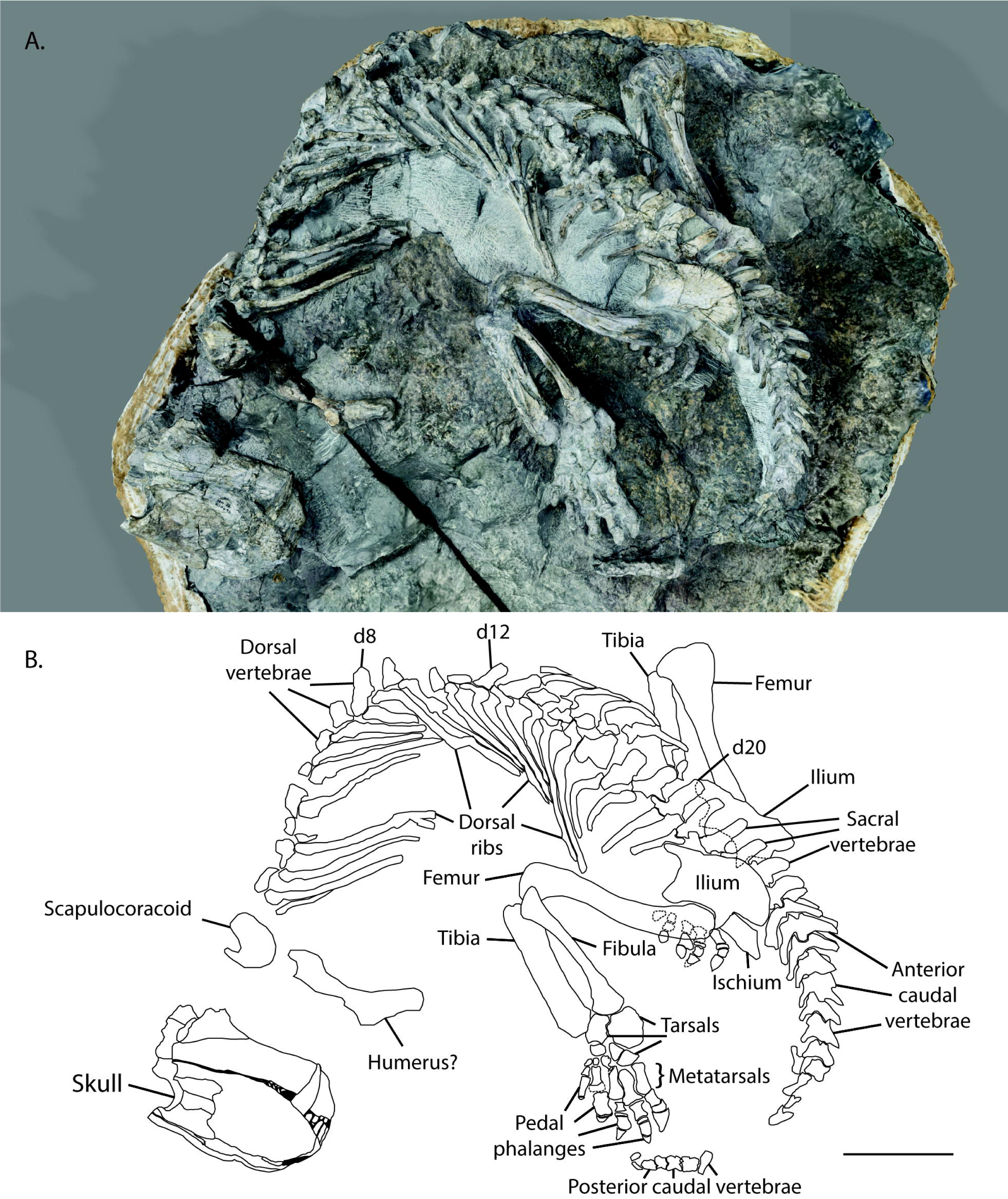
**A photogrammetric 3D render (A) and line drawing (B) of the entirety of *Aelurognathus tigriceps* (SAM-PK-K10496). Scale bar equals ten centimetres.**

SAM-PK-K10496 was excavated in 2004 by Roger M. H. Smith from the farm Doornplaats (cadastral name Rust 126), Graaff-Reinet District, Sarah Baartman District Municipality, Eastern Cape Province, and was prepared by Annelize Crean and Georgina Farrell at the Iziko South African Museum, Cape Town. The specimen was found in the *Dicynodon*-*Theriognathus* Subzone of the *Daptocephalus* Assemblage Zone (Late Permian, 255.2–253.5 Ma (Viglietti *et al*. 2017; Smith *et al*. 2020; Viglietti 2020)) in rocks of the Daggaboersnek Member of the Balfour Formation. Two other gorgonopsian taxa, *Cyonosaurus* and *Rubidgea*, have also been reported from similar horizons at Doornplaats (Viglietti 2020).

### Institutional Abbreviations

AMNH FARB, American Museum of Natural History, Fossil Amphibian, Reptile, and Bird Collection, New York City, NY, USA; BP, Evolutionary Studies Institute (ESI; formerly Bernard Price Institute for Palaeontological Research), University of the Witwatersrand, Johannesburg, South Africa; BSPG, Bayerische Staatssammlung für Paläontologie und Geologie, Munich, Germany; GPIT, Paläontologische Sammlung, Eberhard Karls Universität Tübingen, Tübingen, Germany; NHCC, National Heritage Conservation Commission, Lusaka, Zambia; PIN, Paleontological Institute of the Russian Academy of Sciences, Moscow, Russia; RC, Rubidge Collection, Wellwood, Graaff-Reinet, South Africa; SAM, Iziko South African Museum, Cape Town, South Africa; UMZC, University Museum of Zoology, Cambridge, UK.

## Methods

A 3D photogrammetric scan was made of SAM-PK-K10486 using Abound on iOS (formerly “Metascan”, v3.1.2) (Avrahami et al. 2023). The scan is comprised of 300 photos of the original specimen taken using the main camera of an iPhone 13, processed under the Raw setting with Object Masking enabled, and is available to view online at MorphoSource (https://www.morphosource.org/concern/media/000891666?locale=en). The final scan has a geometry of 3,769,000 triangles. The scan complements separate specimen photos taken with a Panasonic Lumix DMC-FZ300 model camera.

Comparisons with other gorgonopsian specimens were made from the published literature (particularly Broom and Haughton 1913; Colbert 1948; von Huene 1950; Sigogneau-Russell 1989; Tatarinov 2004; Gebauer 2014; Kammerer 2016; Sidor 2022; Bendel et al. 2023; Mann and Sidor 2025), photographs by CFK, and direct examination of the ESI, SAM, and UMZC gorgonopsian collections by SKP.

### Systematic Palaeontology

Synapsida Osborn, 1903

Therapsida Broom, 1905

Gorgonopsia Seeley, 1894

Rubidgeinae Broom, 1938

*Aelurognathus* Haughton 1924

*Aelurognathus tigriceps* (Broom and Haughton 1913)

### Skull

The anterior part of the skull is preserved up to orbital mid-length (Fig. 2). The premaxillae, maxillae, septomaxillae, nasals, prefrontals, lacrimals, anterior half of the frontals, and the anterior portion of the jugals are preserved and visible. The skull is crushed mediolaterally. This is most apparent in dorsal view, as the prefrontals and nasals, which should be symmetrical across the midline, instead have different mediolateral widths (e.g. right prefrontal width = 19 mm vs. left = 11 mm). The ventral surface of the skull is covered by matrix. A transverse cross-section through the orbit exposes part of the pterygoid. The snout is substantially taller dorsoventrally (100 mm from the dorsal margin of the nasal to the ventral margin of the dentary) than it is wide mediolaterally (37.3 mm at the maximum width between the lateral margins of the frontals), although these proportions have probably been exaggerated by crushing. The snout in this specimen is absolutely smaller than in the holotype of *Aelurognathus tigriceps* (SAM-PK-2342) (200 mm dorsoventrally, 101 mm mediolaterally [SKP pers. obs.]), but shows similar proportions, being relatively tall with a convex dorsal margin (Kammerer 2016). The mediolateral width of the skull is approximately consistent throughout the preserved material. The relative width of the snout has taxonomic and phylogenetic importance in Gorgonopsia, especially relative to that of the occiput (Sigogneau 1970; Kammerer 2016). Unfortunately, this index cannot be determined in the current specimen because the postorbital region of the skull is not preserved.

**Fig. 2.**
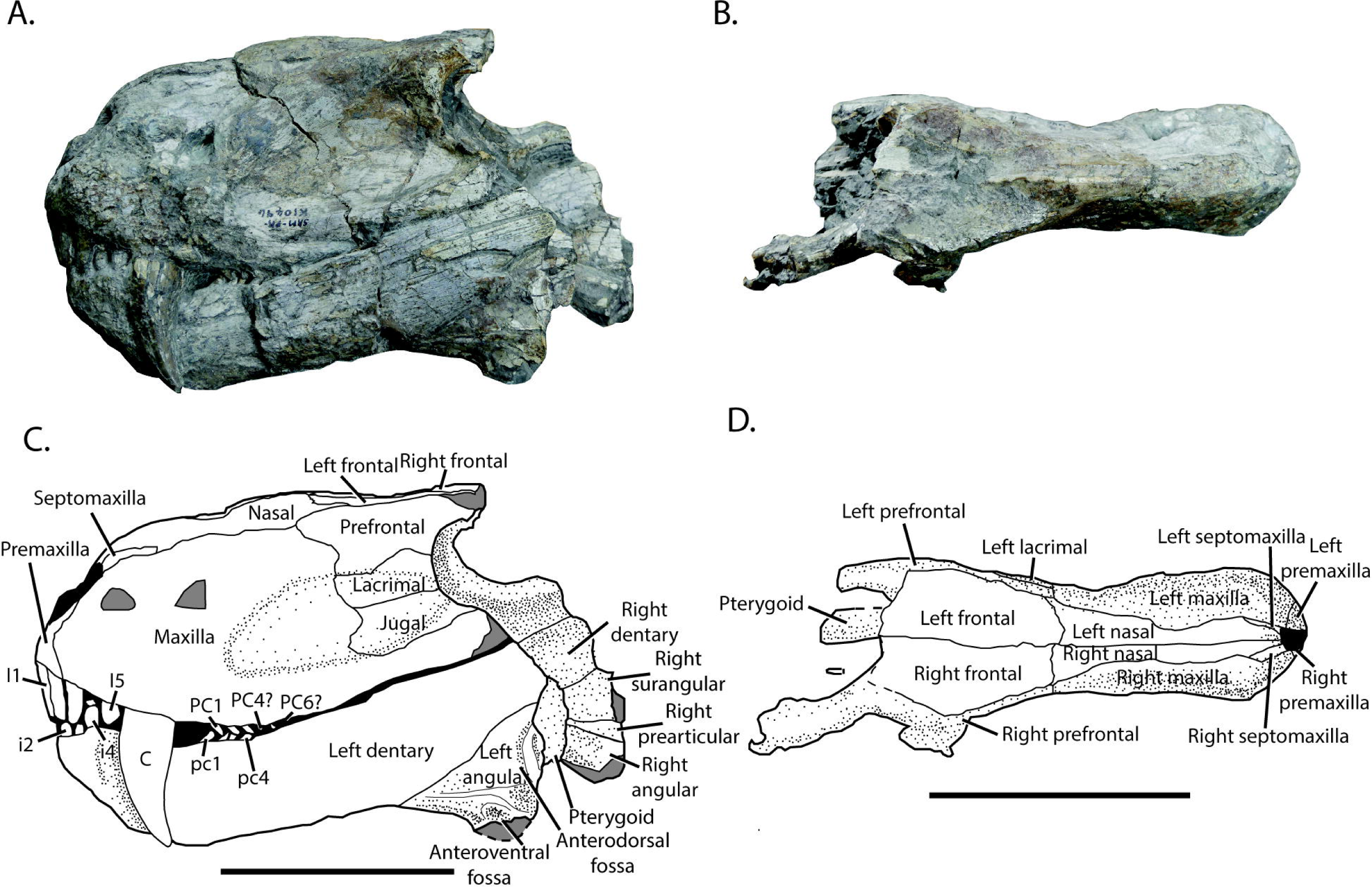
Photographs (A-B) and labelled line drawings (C-D) of the skull of *Aelurognathus tigriceps* (SAM-PK-K10496) from the lateral (A, C) and dorsal (B, D) views. Black areas indicate an opening filled with matrix; light grey areas indicate broken bone. All scale bars equal seven centimetres. Abbreviations: I = upper incisor, i = lower incisor, C = canine, PC = upper postcanine, pc = lower postcanine.

The specimen shares cranial features with other rubidgeines, including the lack of a frontal contribution to the orbit and the tall, robust morphology of the mandibular symphysis (Kammerer 2016). Other important features diagnosing the subclade Rubidgeinae (palatal anatomy, shape of the basicranial girder) are also not visible/preserved in this specimen. Within rubidgeines, the high postcanine count (probably six uppers), weak offset between the facial and zygomatic portions of the maxilla, and lack of pachyostosis (distinguishing it from *Dinogorgon*, in which even the anterodorsal margin of the orbit is notably expanded and rugose) supports identification as *Aelurognathus tigriceps*. The preserved portion of the suborbital zygoma in this specimen is dorsoventrally lower in this specimen than in SAM-PK-2342, although this is within the range of variation within *Aelurognathus,* described by Sigogneau (1970), Gebauer (2007), Norton (2012), and Kammerer (2016).

#### Premaxilla

The left premaxilla is exposed anterolaterally and parts of at least five incisors are visible, although I4 is represented only by a broken root (Fig. 2C). The presence of five incisors is widespread among gorgonopsians (Sigogneau-Russell 1989), except for *Inostrancevia*, which has four (Kammerer *et al*. 2023; Brant & Sidor 2024). The internarial bar is broken off, as is often the case in theriodonts, resulting from its delicate nature (Kammerer 2016). The exposure of the premaxilla itself is minimal, with much of the lateral surface of the premaxilla is covered by the anterior part of the maxilla as in other rubidgeines (Kammerer 2016). The exposed part of the premaxilla is rectangular, with the dorsal margin meeting the external naris and the septomaxilla (Fig. 2C). The posterior margin of the premaxilla is angled posteroventrally below an anterior eminence of the maxilla. The incisors are almost straight, with little curvature, and are proportionally narrow with dorsoventral lengths of approximately 15.3 mm and mesiodistal widths of around 5.9 mm each, with little variation between individual incisors. Apicobasal serrations are not visible on the incisors, although this is likely to be preservational, as the incisors are serrated in most gorgonopsians, including other specimens of *Aelurognathus* (Haughton 1915; Sigogneau 1970). The apices of the incisors are narrower than the basal ends, displaying a slight taper. Their cross-sectional shapes cannot be determined.

#### Maxilla

The left maxilla is the largest of the preserved cranial bones. It has a tall, anteroposteriorly broad dorsal ramus that contacts the nasal dorsally and forms a subvertical contact with the prefrontal, lacrimal, and jugal posteriorly (Fig. 2C). The maxilla continues posteriorly as a tapering process ventral to the jugal, which is broken off below the anterior portion of the orbit. The lateral surface of the maxilla appears rugose due to abrasion, particularly around the canine.

The maxilla preserves a blade-like canine and at least five, possibly six, conical postcanine teeth (Fig. 2C). The canine is 39.5 mm long and 17.0 mm wide at the base of the crown, and its apex extends a few millimetres past the ventral margin of the dentary. The canine displays denticles only along the apical third of the distal carina. However, denticles are likely to have been present originally along the entire distal margin, as is generally the case in rubidgeines (Kammerer 2016). The distal curvature of the canine is slight, approximately 10-15° apicodistally. The mesial edge of the canine also forms a carina but appears to lack denticles. The postcanine teeth are considerably smaller than the other maxillary and premaxillary teeth, with apicobasal lengths up to 3.9 mm and mesiodistal widths of 3.1 mm at the base of each crown. The postcanines show weaker recurvature than the canines, curving between 5-10° apicodistally, but both upper and lower postcanines have a distinct posterior cant. The distal carinae of the postcanines are covered by small amounts of matrix, making it difficult to determine whether denticles are present. The number of postcanines is similar to the type specimen of *Aelurognathus tigriceps* (SAM-PK-2342) as well as a number of other specimens referred to *Aelurognathus* (e.g. BP/1/813, BP/1/1566, BP/1/3464, RC 34, RC 115, SAM-PK-2672, SAM-PK-K10960).

#### Septomaxilla

The left septomaxilla forms most of the posterior edge of the external naris, and its lateral exposure extends posteriorly from the posterodorsal margin of the external naris, between the maxilla and the nasal, to approximately the level of the canine (Fig. 2C). None of the intranarial portion of the septomaxilla is visible; most of its contact with the premaxilla is eroded off. What is preserved of the facial portion approximates a narrow parallelogram. The external surface of the septomaxilla is smooth, lacking the rugosity of the maxilla.

#### Nasal

The nasals are anteroposteriorly long and mediolaterally narrow (Fig. 2D). They contact the external naris anteriorly, and extend posteriorly to contact the prefrontals and frontals. They are subrectangular in dorsal view (with a slight constriction at midlength where the dorsal margins of the maxillae impinge on them), but widen posteriorly.

#### Prefrontal

The prefrontals form the anterodorsal part of the orbital margin and are approximately trapezoidal in lateral view (Fig. 2C). They contact the maxilla and nasal anteriorly, the frontal dorsally, and the lacrimal ventrally. The prefrontal tapers to a point anterodorsally, extending further anterior than the lacrimal or jugal. Along the margin of the orbit, the prefrontals are weakly rugose, showing similar texture to the maxillary surface, but not the pachyostosis and textured ridges seen in, e.g., *Clelandina*, *Dinogorgon* and *Rubidgea* (Kammerer 2016). The posterodorsal edge of the prefrontal is the posteriormost preserved bone in the orbit, so it is uncertain whether the frontal entered the dorsal margin of the orbit. However, based on its transverse thickness at the point of breakage, it is likely that the prefrontal continued to make up the orbital margin further posteriorly, which would exclude the frontal from the orbit if this animal had the usual gorgonopsian morphology in this region (in which the postfrontal makes up the posterior half of the dorsal orbital margin; Kammerer 2016, 2017). Slightly more of the right prefrontal is preserved posteriorly, but its posterior part is obscured by attached matrix.

#### Frontal

The anterior parts of the frontals are preserved and comprise most of the dorsal surface of the cranium as preserved (Fig. 2D). The suture between the frontals and the prefrontals and nasals is visible. The lateral edge of the anterior portion of the frontals is angled anteromedially by approximately 15-20°. On the better preserved left side, the lateral edge of the posterior portion of the frontals is roughly parallel to the long axis of the skull, not showing any indication of angling outwards to contribute to the orbital margin (Fig. 2D).

#### Lacrimal

The left lacrimal is rectangular in lateral view, and composes the central part of the anterior orbit margin, between the prefrontal dorsally and jugal ventrally (Fig. 2C). The lacrimal is proportionally longer anteroposteriorly than it is dorsoventrally. The lacrimal foramina are not visible within the orbit and the facial surface of the lacrimal is too poorly preserved to tell whether an external lacrimal foramen (as in *Dinogorgon* and *Rubidgea*; Kammerer 2016) was present.

#### Jugal

Most of the facial portion of the left jugal is preserved and is approximately rectangular, with a smooth external surface (Fig. 2C). The jugal forms the anteroventral margin of the orbit, contacting the lacrimal dorsally and the maxilla anteriorly and ventrally. In cross-section at the posterior end of the skull, the jugal is wider than the other craniofacial bones, reaching a width of 12.9 mm compared to a cross-sectional width of 8.6 mm for the prefrontal and 7.7 mm for the lacrimal. The preserved extent of the right jugal extends slightly further posteriorly than the left jugal, composing much of the ventral margin of the orbit.

#### Pterygoid

The pterygoid is visible in cross-section only, across the broken posterior surface of the skull as preserved (Fig. 2C-D). Only a small portion of the ventral surface of the transverse processes and palatal boss has been prepared. The cross-section extends from the ventral margin of the skull, past the transverse processes of the pterygoid, and up through the base of the quadrate ramus of the pterygoid, the dorsal section of which is either eroded or obscured by matrix. A small amount of dentition is visible on the ventral boss of the pterygoid. Whereas most of the palatal boss of the pterygoid is obscured by matrix, the visible dentition is comparable to that seen in the *Aelurognathus tigriceps* specimen RC 35 (Kammerer 2016).

### Mandible

In the left mandible, only the dentary and the angular are preserved and exposed, whereas in the right mandible the dentary, angular, surangular, and prearticular are all preserved and at least partially exposed (Fig. 2C). Both dentaries and angulars are broken posteriorly, with slightly more of them preserved on the right than the left.

#### Dentary

Most of the dentary is present, missing only the tip of the coronoid process posterior to the orbital midpoint, although the anterior end is somewhat eroded (Fig. 2C). Its anterior edge is angled anterodorsally at 50-60° above horizontal, and the posterior quarter of the ventral edge is angled 20-30° above the horizontal to border the angular dorsally. The dentary tapers posteriorly to around half of its maximum dorsoventral height at its posteriormost preserved part. As is the case in most gorgonopsians, there is a fossa on the lateral surface of the dentary for the upper canine, and preceding this is a thickening of the bone around the root of the lower canine within the dentary (Fig. 2C).

Four dentary incisors are present, with i1 obscured by an upper incisor, i2 and i3 mostly obscured by upper incisors, but visible at base, and i4 completely exposed (Fig. 2C). At least four postcanine teeth are visible and are oriented posterodorsally, with the distalmost tooth slightly obscured by an upper postcanine; this suggests that there are likely fewer lower postcanines than upper postcanines present in SAM-PK-K1046, as in other rubidgeines (Kammerer 2016). The lower incisors are smaller than the upper incisors but share a similar conical shape.

#### Angular

Only the anterior part of the angular is preserved, with a small section of the reflected lamina present on both the left and right angulars (Fig. 2C). The preserved part of the angular is roughly triangular in shape, with the reflected lamina displaying the horizontal anteroposteriorly-oriented ridge and a section of the strong vertical dorsoventrally oriented ridge, which together bound the anterodorsal fossa (Fig. 2C), as seen in *Cynariops robustus* (MB.R.999) (Olroyd and Sidor 2022). Ventral to the horizontal ridge is a second, smaller ridge that curves posteroventrally and together the two ridges bound the anteroventral fossa (Fig. 2C).

#### Medial mandible bones

A small portion of the medial surface of the right mandible is exposed, showing parts of the dentary, surangular, prearticular, and angular (Fig. 2A, C). The morphology of these bones is comparable to that of other gorgonopsians, such as “*Scymnognathus*” *parringtoni* (GPIT-PV-31579), *Leontosaurus vanderhorsti* (BP/1/803), and *Arctops umulunshi* (NHCC LB396) (von Huene 1950; Sigogneau 1970; Kammerer 2016; Mann and Sidor 2025). The surfaces of all the medial mandible bones are smooth, and show slightly different angulation. The dorsal margin of the dentary is angled very slightly (5-10°) dorsally, and the surangular is similarly angled. However, the prearticular and angular are angled more ventrally (∼15-20°).

### Dorsal vertebrae

The cervical vertebrae are missing and it is difficult to determine the identity of the anteriormost preserved dorsal vertebra. However, it is clear that most of the dorsal vertebral column is preserved, with twenty-one dorsal vertebrae present, numbered here d1-d21 from anterior to posterior (Fig. 1B). All centra are obscured, either by matrix or the dorsal ribs. Many neural spines are broken or missing, other than in d8, d12, d13, d16, d18, d19, d20, and d21. The transverse processes are approximately square in dorsal view, with the mediolateral length just slightly greater than the anteroposterior width, and dorsoventrally thin (∼1.5 mm). The transverse processes are generally short (e.g. 16.0 mm in d6) relative to the vertebral length from the anteriormost tip of the prezygapophyses to the posteriormost tip of the postzygapophyses (e.g. 31.1 mm in d6; henceforth: ‘zygapophyseal vertebral length’).

The pedicles are mediolaterally short, with little distance between the zygapophyses and transverse processes. While the left zygapophyses are exposed for most dorsal vertebrae, the right zygapophyses are all entirely covered by matrix. The prezygapophyses compose roughly a third of the total zygapophyseal vertebral length. They are angled nearly vertically, roughly 5-10° from the sagittal plane. Their articular surfaces are approximately oval, with the major axis being roughly double the length of the minor axis. The postzygapophyses also compose roughly a third of the zygapophyseal vertebral length. However, the angle of the postzygapophyses is far less sharp than that of the prezygapophyses, being closer to 45° from the anteroposterior midline.

The neural spines are mediolaterally narrow and are angled >45° posteriorly (Fig. 1B). The dorsal neural spines increase in size posteriorly; d20 has the largest neural spine, with an anteroposterior length ∼12.9 mm, and dorsoventral height of 27.4 mm. The neural spines of the anteriormost preserved dorsal vertebrae are missing, meaning it is not possible to measure their lengths, but d1 has a mediolateral width of 24.3 mm, compared to 15.2 mm for d18, the posteriormost dorsal vertebra for which this measurement can be taken (d19-d21 are covered by matrix, preventing measurement).

### Dorsal ribs

Dorsal ribs are present in articulation throughout the dorsal series, with left ribs preserved and exposed for every dorsal vertebra from d1 through to d21 and right ribs preserved and exposed for d5-d8 (although the ribs for d5 and d6 are posteriorly displaced), d13-d15, d17, d18, d20, and d21 (Fig. 1B). The posteriormost three ribs (d19-d21) have a distinct lumbar morphology, differentiated from the preceding thoracic ribs, with d18 displaying a transitional morphology. The thoracic ribs are all bicapitate, whereas the lumbar ribs display only one head. This single-head condition of the lumbar ribs is hypothesized to have been present in other gorgonopsian specimens, such as “*Aelurognathus*” *microdon* (SAM-PK-9344) (Boonstra 1934), *Lycaenops ornatus* (AMNH FARB 2240) (Colbert 1948), and “*Scymnognathus*” *parringtoni* (GPIT-PV-31579) (Gebauer 2014), but in SAM-PK-K10496 it is directly observable. The thoracic ribs are elongate and gently curve posteroventromedially. By contrast, the lumbar ribs are differently angled, curving more anteroventrally, and are far shorter in length than the thoracic ribs (Fig. 1). The degree of medial curvature of the left ribs is obscured by mediolateral crushing of the specimen. The right dorsal ribs exhibit greater medial curvature as preserved.

Thoracic ribs are around 10-11 cm long, with the anteriormost preserved rib at ∼10.3 cm and the longest preserved rib at ∼10.8 cm. The lumbar ribs are shorter than the thoracic ribs, with the longest lumbar rib being ∼5.4 cm long and the shortest lumbar rib – also the final dorsal rib – being ∼4.4 cm long (Fig. 1B). The cross-sectional diameters narrow from dorsal to ventral, with the dorsal section being about twice the width of the ventral section. The posteriormost ribs are angled slightly anteriorly, but this may be due to the curvature of the vertebral column in death or displacement from the pelvis in the case of the final dorsal rib, as the final dorsal ribs are positioned ventral to the anterodorsal process of the ilium. The tubercula and capitula are generally obscured by the transverse processes and pedicles of the dorsal vertebrae to which they articulate. In d3, one of the only ribs with an exposed tuberculum and capitulum, the tuberculum and capitulum are closely spaced together, and are roughly the same width as each other.

### Sacral vertebrae

The sacrum consists of three sacral vertebrae (Fig. 1B). However, most of its morphology is obscured by matrix, so it is not possible to determine whether the sacral vertebrae are fused. Only the neural spines and left zygapophyses of the anterior two sacral vertebrae are exposed and only the neural spine, left and right zygapophyses, and transverse processes of the third sacral vertebra are exposed. The transverse processes of the third sacral vertebra are generally rectangular and are longer than those of the dorsal vertebrae. Although their exposed maximum mediolateral length (15.8 mm) is similar to that of the dorsals, the medial section of the right transverse process and the lateral section of the left transverse process are both covered by matrix. The zygapophyseal vertebral length of the third sacral vertebra is 31.3 mm, making the length of the transverse processes relative to the length of the entire vertebra (>1:2) longer than that of the dorsal vertebrae (1:2). The distal ends of the transverse processes are fused to the ilium. The anteroposterior width of the transverse processes is 8.3 mm. The zygapophyses share a similar overall shape to those of the dorsal vertebrae, similar angulation from the anteroposterior midline, and comparable proportions relative to total vertebral length (∼1/3). The neural spines are dorsoventrally taller than those of the dorsal vertebrae, reaching a maximum length of 29.8 mm in the anteriormost sacral vertebra. The longest anteroposterior length for a sacral neural spine is 13.5 mm, occurring in the second sacral vertebra. The neural spines are mediolaterally thin, being no more than 2.5 mm wide. The neural spine is nearly vertically angled in the anterior two sacral vertebrae and only slightly posterodorsally angled in the third.

### Caudal vertebrae

Almost the whole tail is preserved (Fig. 3), which is rare among gorgonopsians (but see Colbert [1948], Tatarinov [2004], and Bendel *et al*. [2023] for prior descriptions of gorgonopsian caudal vertebrae). The tail is broken into two sections, arranged in a continuous curve, with the intervening elements lost to erosion. The first section articulates with the sacrum and includes ca1–ca13, although only chevrons remain for ca11–ca13. The second section comprises ca17–ca23 (estimated identities; Fig. 1B, 3B). At least four caudals are estimated to be missing, based on the difference in size between ca13 and the first vertebra of the second section (ca17) and this minimal number of missing vertebrae is used for our numbering. The maximal number of missing caudals is six, based on dividing the missing length by the anteroposterior diameter of the posterior vertebrae of the anterior section or the anterior vertebrae of the posterior section. The terminal two preserved caudals (here numbered c22 and c23) are simple, disc-like structures as exposed, unlike the more elongate terminal caudal vertebrae seen in theriodonts with definite complete caudal series (e.g. Butler *et al*. 2019). Based on this, there could be one or two vertebrae missing at the end of the tail, making the total number of caudal vertebrae between 23 and 27.

**Fig. 3.**
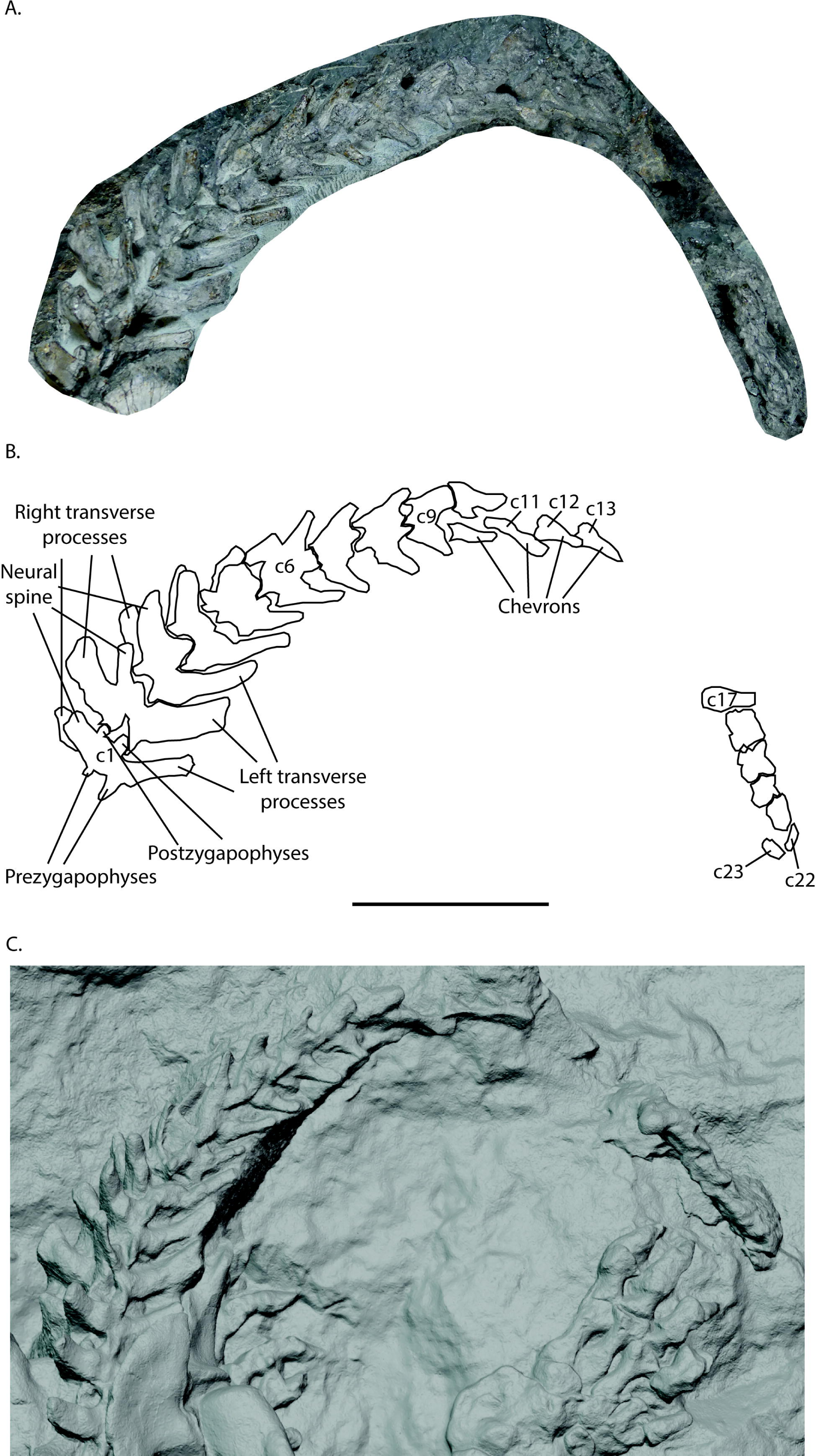
A photograph (A), labelled line drawing (B), and rendering of a photogrammetric model (C) of the tail of *Aelurognathus tigriceps* (SAM-PK-K10496) from the dorsolateral view. The scale bar is eight centimetres long.

The caudal centra are obscured by matrix, with only the transverse processes, zygapophyses, and neural spines exposed. Chevrons are exposed for ca10–ca13 (Fig. 3B). Left transverse processes are visible for ca1–ca9 and right transverse processes are visible for ca1–ca6. Zygapophyses are preserved in all caudal vertebrae except ca11–ca13. Neural spines are present for ca1-ca10, and the absence of neural spines in ca17–ca23 is likely real morphology and not due to preservation (Fig. 3B). Anterior vertebrae have larger transverse processes, with a maximum mediolateral length of 35.8 mm in ca2. This is proportionally longer than in any other section of the vertebral column, as the zygapophyseal vertebral length of ca2 is 31.7 mm (Table 1), thus exceeding a 1:1 ratio. This large proportion begins to diminish around ca7 and ca8, with the transverse processes sharply reducing in mediolateral length until ca10 (Table 1). Transverse processes are absent in ca17 and more posterior caudals. In ca1–ca6, the transverse processes are angled sharply posteriorly, at 45-60° from the mediolateral horizontal midline. The zygapophyses are similarly shaped to those elsewhere in the vertebral column (Fig. 1B, 3B). In the anterior section of the tail, the ratio of pre- and postzygapophysis diameter to zygapophyseal vertebral length is the same as is seen in the dorsal and sacral vertebrae (1:3), whereas for the tail tip it is closer to 1:4. The angle of the prezygapophyses is less vertical than elsewhere in the vertebral column, being closer to 30° lateral to the sagittal plane (Fig. 3B). The neural spines are tall in the anterior caudal vertebrae, reaching a maximum dorsoventral length of 24.6 mm in ca1, and gradually decreasing posteriorly (Table 1). The neural spines also begin to be angled more posterodorsally with each posterior vertebra. The neural spines reach a minimum dorsoventral length of 14.8 mm in ca10, before being lost entirely in ca17–ca23. The chevrons seen in ca10–ca13 are long, with proximodistal lengths of ∼23.5 mm. The chevrons sharply narrow as they extend posteriorly, with proximal mediolateral widths of ∼11.5 mm and distal mediolateral widths of ∼4.0 mm (Table 1). They also display a flat blade at their distal tips, with a dorsoventral height of ∼8.1 mm. The ca17–ca23 are proportionally longer than the more anterior caudal vertebrae, indicating a slight elongation in posterior caudal vertebrae similar to that seen in *Viatkogorgon ivakhnenkoi* (PIN 2212/61) (Tatarinov 2004). The presence of different proportions between anterior and posterior caudal vertebrae in *Viatkogorgon ivakhnenkoi* led Tatarinov (2004) to suggest a semi-aquatic locomotor mode. However, the presence of similar morphologies here in SAM-PK-K10496 suggests that this may be characteristic of gorgonopsian tails generally, and not a specific indication of aquatic habits, given the lack of any other indicators that SAM-PK-K10496 has distinct semi-aquatic adaptations.

**Table 1.**
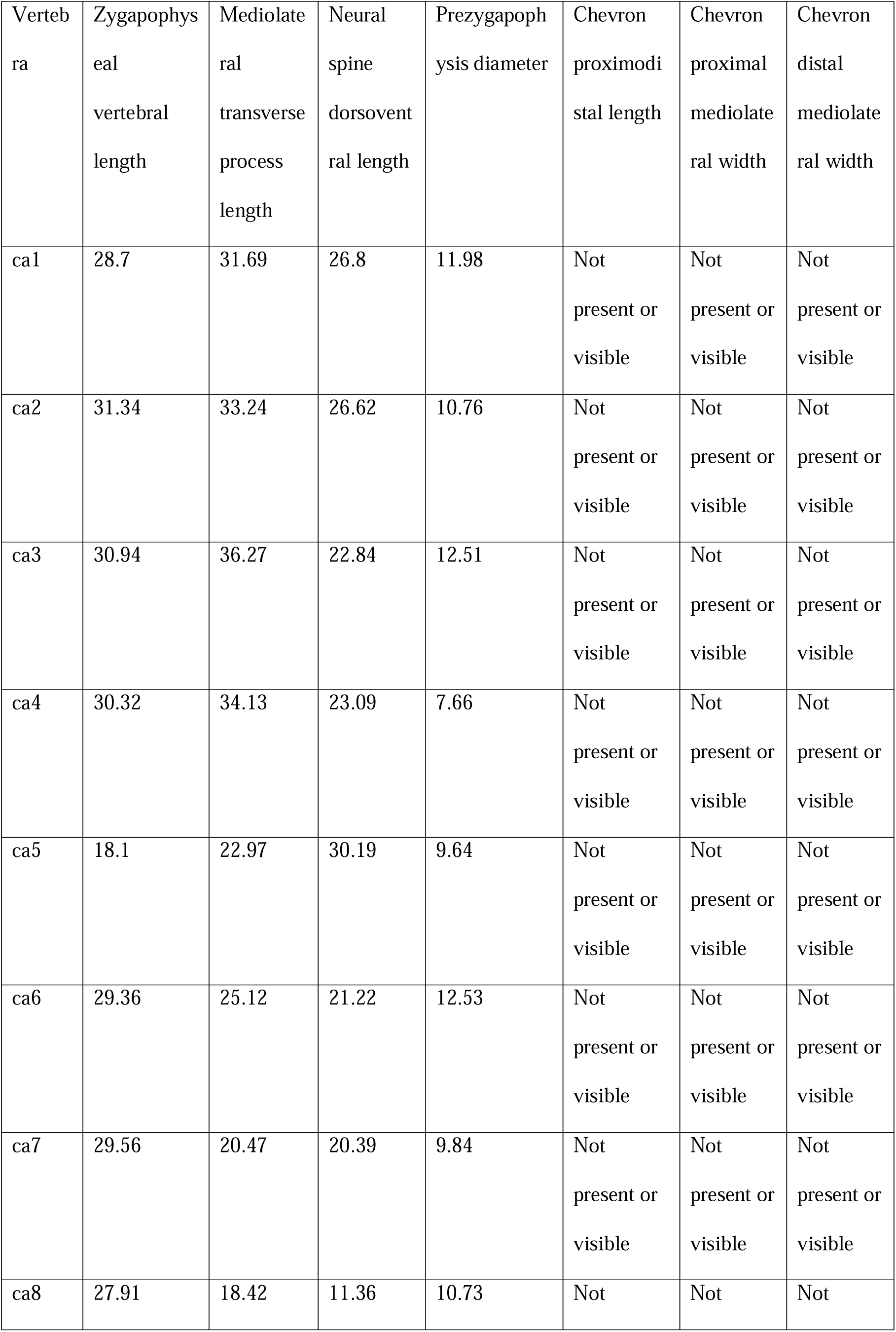
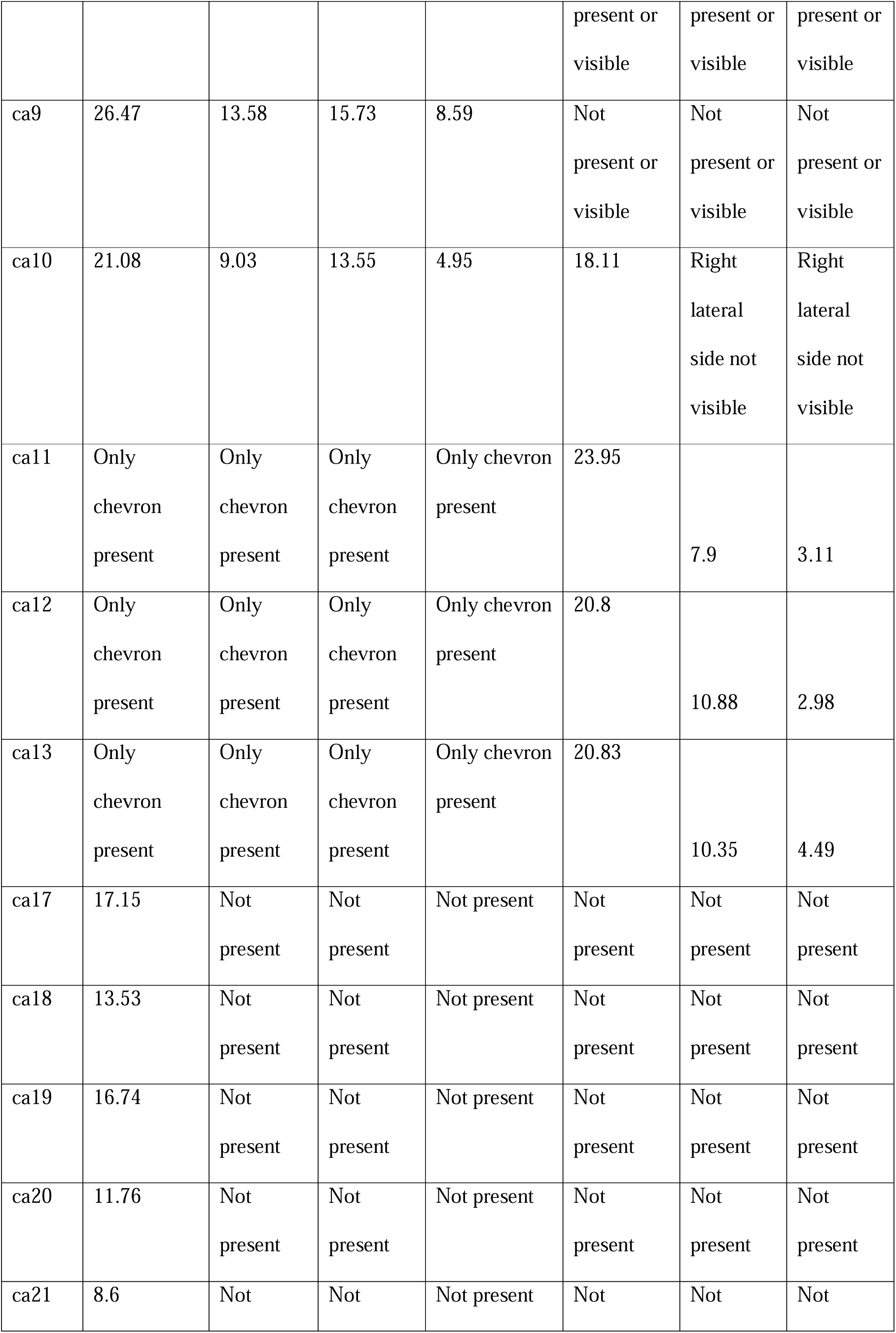
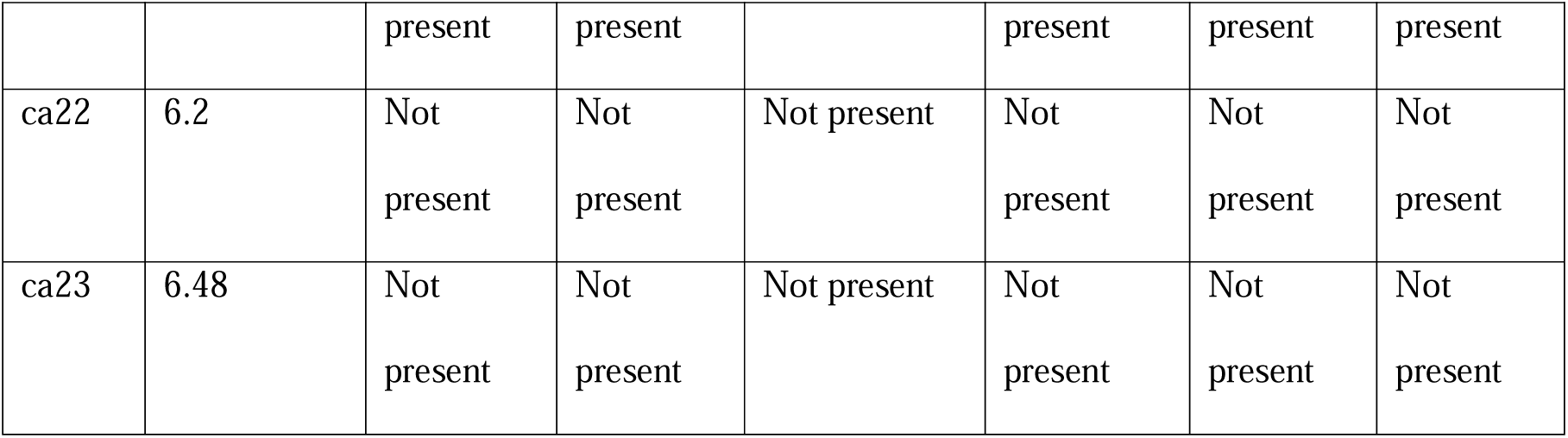
Length and width measurements for the morphology of the caudal vertebrae of *Aelurognathus tigriceps* (SAM-PK-K10496). All measurements are listed in millimetres.

### Pectoral girdle

A partial left scapulocoracoid is preserved in SAM-PK-K10496 (Fig. 4). Much of the surface of the scapulocoracoid is weathered and broken, making precise identification of the preserved elements difficult. The identifiable exposure includes the ventral portion of the scapular blade, and likely the coracoid and procoracoid (Fig. 4B). The scapular blade seems mediolaterally narrow compared to those of *Aelurognathus tigriceps* (SAM-PK-2342) or *Lycaenops ornatus* (AMNH FARB 2240) instead of broad as in *Gorgonops torvus* (SAM-PK-K10591) (Broom and Haughton 1913; Colbert 1948; Bendel et al. 2023). However, this could be a result of the lateral taphonomic compression endured by the whole specimen. Ascertaining the precise morphology of the preserved material in the ventral half of the preserved scapulocoracoid is difficult due to the heavy erosion and presence of matrix surrounding the bone.

**Fig. 4.**
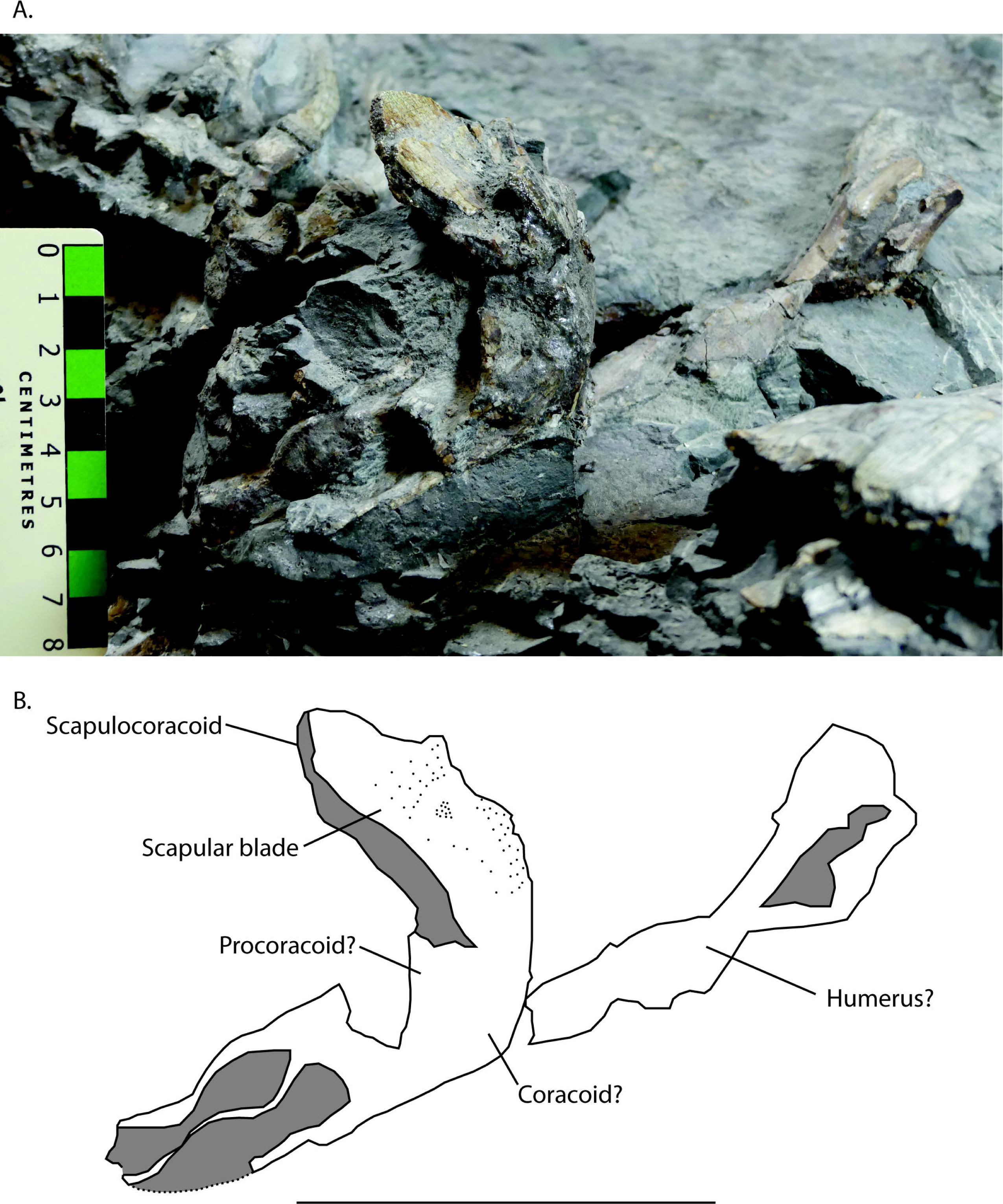
A photograph (A) and labelled line drawing (B) of the left pectoral girdle and possible forelimb of *Aelurognathus tigriceps* (SAM-PK-K10496) from the anterior view. The scale bar is eight centimetres long.

There is also a fragmentary, elongate bone associated ventrally with the left scapulocoracoid, which may be a humerus (Fig. 4B). However, the identity of this element is uncertain.

### Pelvic girdle

The left and right sides of the pelvis are both preserved and articulated, although not all parts are exposed and some parts, including the acetabulum and most of the left pubis, are obscured by the left femur (Fig. 5). The left ilium is fully exposed, while the left ischium is only partially exposed (Fig. 5C), and the left pubis is mostly obscured, with its ventral surface exposed underneath and medial to the left femur (Fig. 5D). On the right, only the ilium is visible. The iliac blade has a convex dorsal margin, giving the ilium a curved dorsal outline in lateral view, with a rounded posterodorsal corner and a prominent anterodorsal process. The iliac blade is mediolaterally thin and anteroposteriorly broad, with an anteroposterior length of ∼96 mm. The iliac contribution to the acetabulum is obscured by the femur. The anterodorsal process of the left ilium extends ∼20 mm beyond the pubic articulation, extending dorsal to the posteriormost dorsal rib, and is dorsoventrally narrow, comprising 20% of the length of the anterior edge of the ilium (Fig. 5C). The anterodorsal process of the ilium of SAM-PK-K10496 is therefore proportionally longer (Table S1) than in most other gorgonopsians, including unidentified gorgonopsian UMZC T883 (Table S1 [SKP pers. obs.]), *Lycaenops ornatus* (AMNH FARB 2240; [Table S1]), *Gorgonops torvus* (SAM-PK-K10591; [Table S1]), and *“Scymnognathus” parringtoni* (GPIT-PV-31579; [Table S1]). It is comparable in proportional size only to *“Aelurognathus” microdon* (SAM-PK-9344; [Table S1]) among published gorgonopsian specimens (Boonstra 1934; Colbert 1948; Gebauer 2014; Bendel et al. 2023). The dorsoventrally narrow proportions of the anterodorsal process of SAM-PK-K10496 (20% the dorsoventral length of the anterior edge of the ilium [Table S1]) are closer to those of “*Scymnognathus*” *parringtoni* (GPIT-PV-31579; [Table S1]) than those of UMZC T883 (Table S1 [SKP pers. obs.]), “*Aelurognathus*” *microdon* (SAM-PK-9344; [Table S1]), and *Lycaenops ornatus* (AMNH FARB 2240; [Table S1]), which comprise a greater proportion of the anterior edge of the ilium (Boonstra 1934; Colbert 1948; Gebauer 2014). The angle of curvature from the anterodorsal process to the anterior edge between the iliac crest and the pubis is approximately 126° in SAM-PK-K10496.

**Fig. 5.**
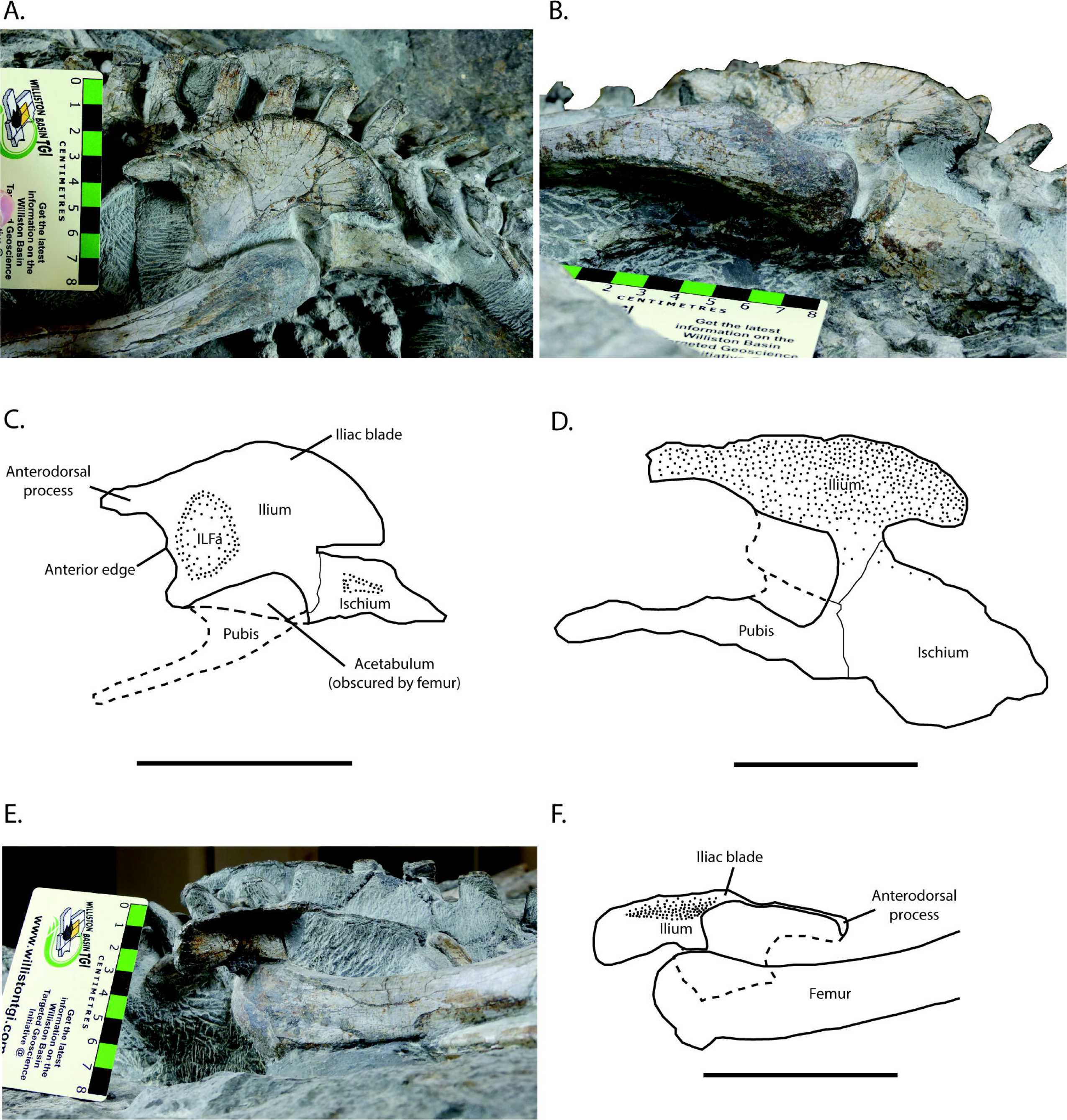
Photographs (A-B, E) and labelled line drawings (C-D, F) of the pelvic girdle of *Aelurognathus tigriceps* (SAM-PK-K10496) from the left lateral (A, C), ventral (B, D), and right lateral (E-F) views. Scale bar for both lateral views is eight centimetres long, and the scale bar for the ventral view is five centimetres long. ILFa stands for the anterior head of the iliofemoralis muscle.

The portion of the iliac blade of SAM-PK-K10496 that dorsally overhangs the ischium (24 mm, Table S1) is proportionally similar to that of UMZC T883 (30 mm, Table S1 [SKP pers. obs.]) and larger than that of *Lycaenops ornatus* (AMNH FARB 2240 [15 mm, Table S1 [SKP pers. obs.]) (Colbert 1948; Bendel et al. 2023). There is a shallow depression on the anterolateral surface of the left ilium which represents the origin of the anterior head of the iliofemoralis muscle (Bishop and Pierce 2024) (Fig. 5C). As preserved, the left and right ilia show differing degrees of mediolateral curvature, with the left ilium being approximately flat and the right curving anteroposteriorly so the lateral surface is concave (Fig. 5E-F).

The ischium is dorsoventrally tall anteriorly, and tapers posteriorly from approximately its midlength (Fig. 5C). The ischium is incomplete, with the posterior half broken and eroded, which contributes to its smaller apparent size when compared to other specimens, such as indeterminate gorgonopsian UMZC T883 (SKP pers. obs.), *Lycaenops ornatus* (AMNH FARB 2240), and *Gorgonops torvus* (SAM-PK-K10591) (Colbert 1948; Bendel et al. 2023). The dorsal and ventral edges of the anterior half of the ischium appear to parallel each other (Fig. 5C). The ischium extends directly posteriorly, as seen in “*Aelurognathus*” *microdon* (SAM-PK-9344), rather than posteroventrally at a 45° angle as in *Lycaenops ornatus* (AMNH FARB 2240), unidentified gorgonopsian UMZC T883 (SKP pers. obs.), and “*Scymnognathus*” *parringtoni* (GPIT-PV-31579) (Colbert 1948; von Huene 1950; Sigogneau 1970; Gebauer 2014; Bendel et al. 2023).

The pubis is rod-like, with the exposed section being far narrower dorsoventrally than either the ilium or ischium (Fig. 5D). Near the contact of the pubis and ischium there appears to be some breakage or erosion of the bone, making the exact contact between the two bones difficult to identify. This state of preservation also makes it difficult to identify the possible presence of foramina, such as the pubic foramen or the obturator foramen. The shape of the pubis in SAM-PK-K10496 is similar in its mediolateral narrowness as viewed ventrally to that of unidentified gorgonopsian UMZC T883 and *Gorgonops torvus* (SAM-PK-K10591) as opposed to how the pubis broadens distally and medially in *Lycaenops ornatus* (AMNH FARB 2240); it is possible that the latter condition is present in SAM-PK-K10496, but this cannot be confirmed because of obstruction by matrix (Colbert 1948; Bendel et al. 2023).

### Femur

Both femora are well preserved and exposed, although the femoral heads are obscured by both matrix and the articulated pelvic bones (Fig. 5D-E, 6, 7A-C). The distal condyles of the right femur are clearly exposed, whereas the medial condyle of the left femur is obscured by matrix. The femur overall has a slight S-curve in lateral view, with the proximal end curving slightly anteroproximally, and the distal end curving slightly posterodistally (Fig. 6B). In anterior view, the proximal end of the femur curves medially 20-30° from the proximodistal midline, whereas the distal end curves laterally 5-10° from the proximodistal midline (Fig. 7B). The femoral shaft is proportionally slender (Table S1), similar to some other gorgonopsians such as *Cyonosaurus* sp. (SAM-PK-K10539) (Table S1 [SKP pers. obs.]) and indeterminate gorgonopsian SAM-PK-K11207 (Table S1 [SKP pers. obs.]), while being less slender than *Gorgonops torvus* (SAM-PK-K10591) (Table S1 [SKP pers. obs.]), and more slender than *Lycaenops ornatus* (AMNH FARB 2240) (Table S1 [SKP pers. obs.]), “*Aelurognathus*” *microdon* (SAM-PK-9344) (Table S1 [SKP pers. obs.]), indeterminate gorgonopsian SAM-PK-K10585 (Table S1 [SKP pers. obs.]), and *Gorgonops* sp. (BSPG-1934 VIII 28) (Table S1 [SKP pers. obs.]) (Boonstra 1934; Broili and Schröder 1935; Colbert 1948; Bendel et al. 2023). The mediolateral width of the proximal end is 26.5 mm, smaller than the mediolateral width of the distal end, of 50.1 mm, and both widths are larger than the midshaft mediolateral width of 20.6 mm.

**Fig. 6.**
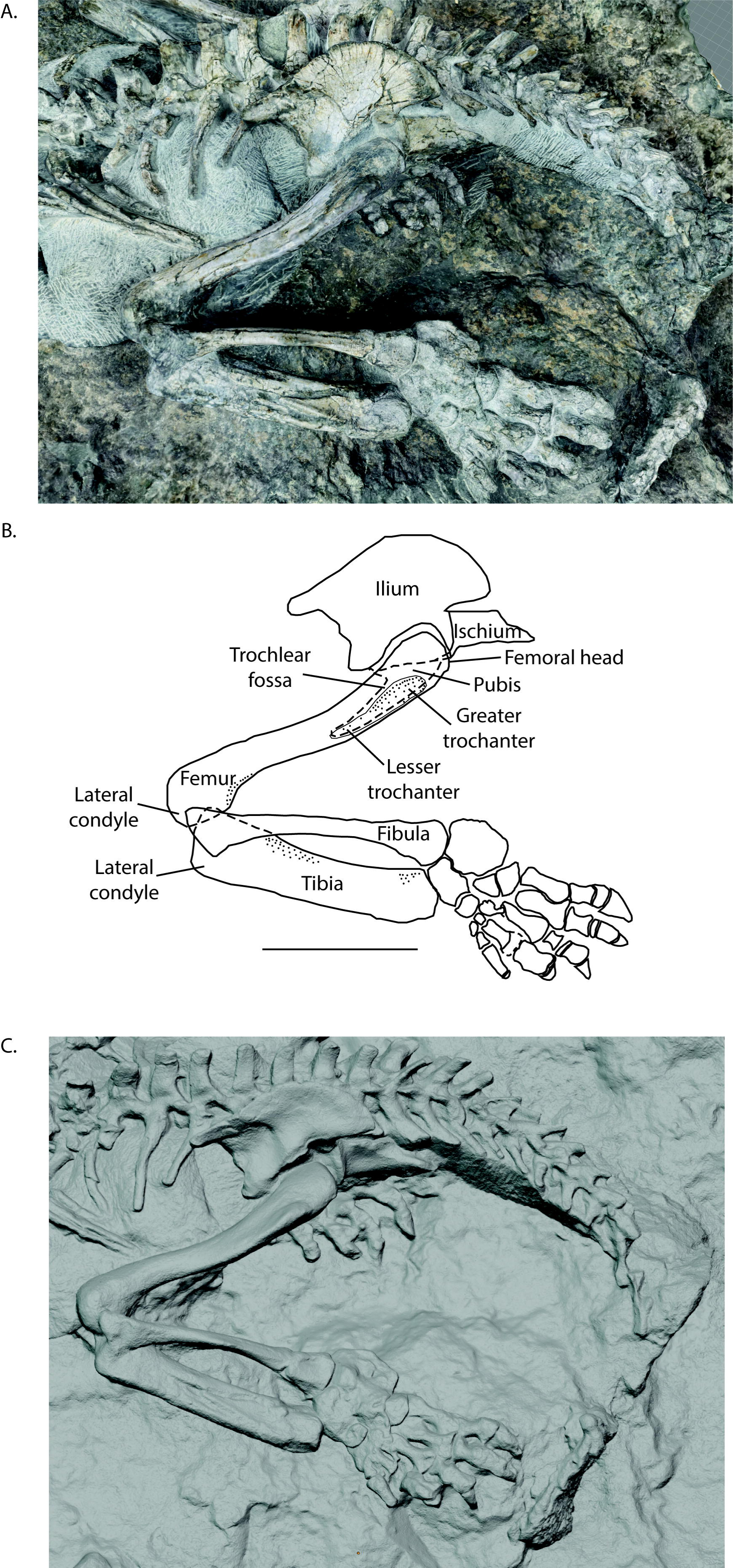
A textured photogrammetric 3D render (A), labelled line drawing (B), and texture-free rendering of a photogrammetric model (C) of the left pelvic girdle, hind limb, and pes of *Aelurognathus tigriceps* (SAM-PK-K10496). The scale bar is eight centimetres long.

**Fig. 7.**
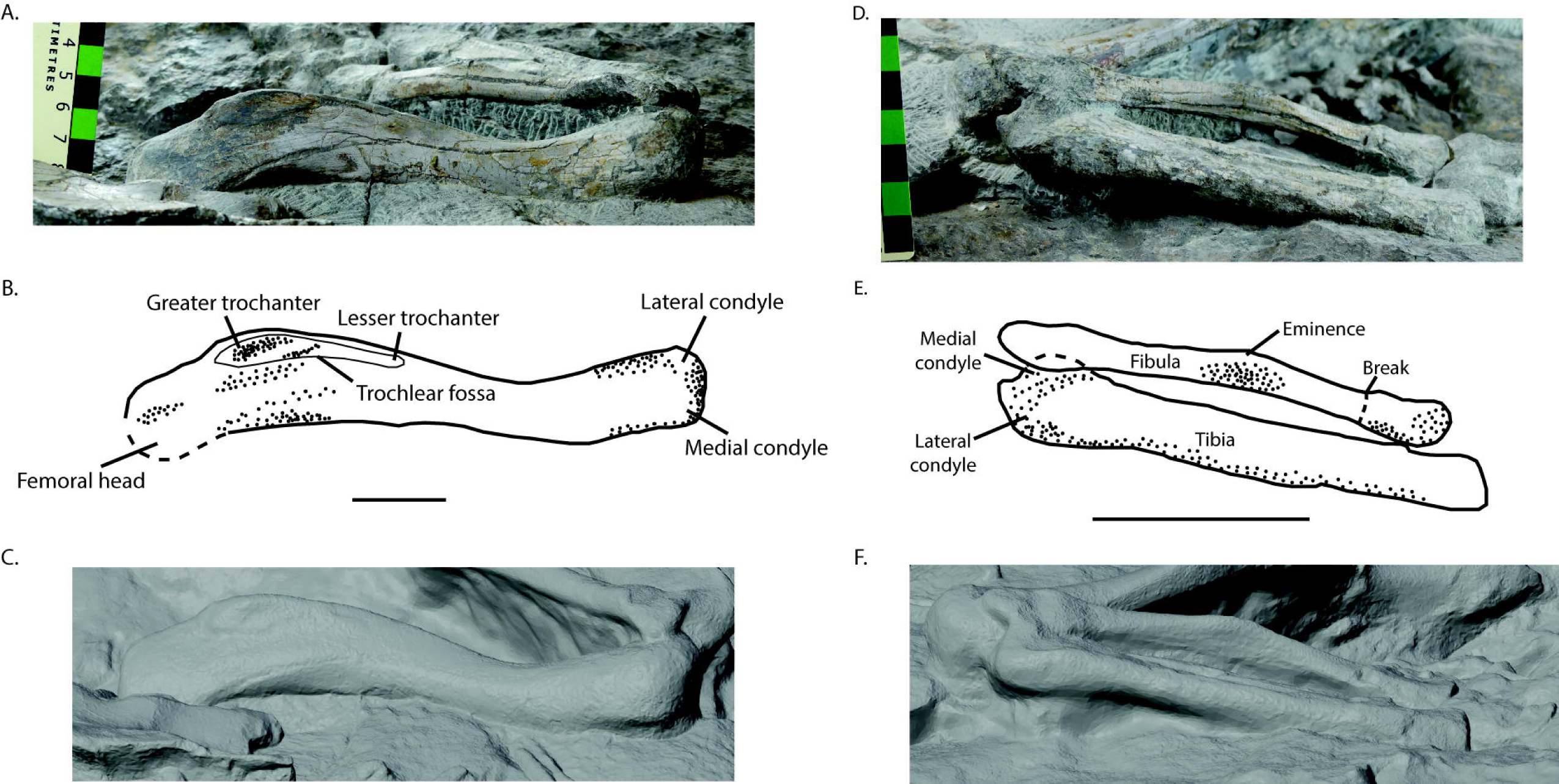
Photographs (A, D), labelled line drawings (B, E), and renderings of a photogrammetric model (C, F) of the left femur (A-C) & left tibia and fibula (D-F) of *Aelurognathus tigriceps* (SAM-PK-K10496). All images are shown in anterior view. The scale bar for the femur is three centimetres and the scale bar for the tibia and fibula is six centimetres.

The greater trochanter is prominent and is located on the lateral surface about two centimetres distally from the proximal end of the femur (Fig. 7B). The left femur has a maximum proximodistal length of ∼169.9 mm, and the greater trochanter is ∼42.5 mm long, therefore occupying slightly more than one-fifth of the overall femur length. The greater trochanter also has a mediolateral depth of 10.7 mm, which is about one third of the total width of the femur at that location of 29.1 mm. The lesser trochanter is greatly reduced compared to other gorgonopsian specimens, such as *Gorgonops torvus* (SAM-PK-K10591), *Lycaenops ornatus* (AMNH FARB 2240), *Gorgonops* sp. (BSPG 1934 VIII 28) and unidentified gorgonopsian SAM-PK-K10585 (SKP pers. obs.) where it is larger and more readily identifiable, instead being more similar in size to that of “*Aelurognathus*” *microdon* (SAM-PK-9344) and unidentified gorgonopsian SAM-PK-K11207 (Boonstra 1934; Colbert 1948; Bendel et al. 2023). The distal condyles are separated by a small depression, and have similar sizes to each other, although the lateral condyle is slightly larger in mediolateral width than the medial condyle (Fig. 7B).

### Tibia

The left tibia is exposed and the right tibia is partially exposed but mostly obscured by the right femur and dorsal ribs, and its distal end is embedded in matrix ventral to the sacrum (Fig. 6, 7D-F). Both tibiae are complete, although the left tibia is shattered. The tibia is proportionally more robust than the fibula. The left tibia is dorsoventrally flat, likely due to the taphonomic dorsoventral crushing indicated by the presence of substantial cracks and breaks (Fig. 7E). The proximal end of the tibia has a mediolateral width of 29.2 mm, which is only slightly larger than the mediolateral width of the distal end, at 26.6 mm and the midshaft width of 25.3 mm. The tibia has a proximodistal length of 140.4 mm, making it shorter than the femur and nearly identical in length to the fibula. The tibia overall curves very slightly medially, similar to “*Aelurognathus*” *microdon* (SAM-PK-9344), *Gorgonops torvus* (SAM-PK-K10591), and *Gorgonops* sp. (BSPG 1934 VIII 28) (Broili and Schröder 1935; Sigogneau-Russell 1989; Bendel et al. 2023). The articular surface of the distal end is very slightly concave medially, where it contacts the astragalus (Fig. 6B-C).

### Fibula

The left fibula is fully preserved and exposed, whereas the right fibula is preserved but mostly obscured by the right femur (Fig. 6B, 7D-F). The fibula has a cylindrical shaft with mediolaterally-expanded proximal and distal ends. The fibula has a proximodistal length of 140.4 mm, and a minimum mediolateral shaft width of 9.1 mm. The proximal end has a mediolateral width of 28.9 mm and the distal end has a mediolateral width of 30.5 mm. The fibula bows laterally (Fig. 6B, 7E), similar to *Lycaenops ornatus* (AMNH FARB 2240) (Colbert 1948). This curvature, along with the curvature of the tibia, results in the expanded interosseous space typical of gorgonopsians, differing from other therapsid groups in which the space between the tibia and fibula is lesser (Colbert 1948; Matamales-Andreu et al. 2024). The distal end is broken off from the rest of the fibula, offsetting it slightly mediolaterally (Fig. 7E). The articular surface of the distal end contacts both the calcaneum and the astragalus. A broad eminence is present on the lateral surface of the fibula, around midlength (Fig. 7E), as also seen prominently in *Gorgonops* sp. (BSPG 1934 VIII 28), *Gorgonops torvus* (SAM-PK-K10591), “*Aelurognathus*” *microdon* (SAM-PK-9344), and *Arctops umulunshi* (NHCC LB396) where it is described as a rugosity, but not in *Lycaenops ornatus* (AMNH FARB 2240) (Boonstra 1934; Broili and Schröder 1935; Colbert 1948; Bendel et al. 2023; Mann and Sidor 2025). This eminence may be similar to the fibular crest noted in *Viatkogorgon ivakhnenkoi* (PIN 2212/61), however in SAM-PK-K10496 the eminence does not span much of the length of the fibula as is figured for PIN 2212/61 (Tatarinov 2004).

### Tarsals

The left tarsals are preserved and exposed in dorsal (extensor) view, whereas the right tarsals are likely preserved but are completely obscured by the sacrum and matrix (Fig. 8). The calcaneum and astragalus are present and unobscured at the dorsal surface. The centrale and the cuboid (which in gorgonopsians is formed by the fused distal tarsals (Dt) 4 and 5 (Sigogneau 1970; Sigogneau-Russell 1989; Hopson 1995)) are easily identifiable (Fig. 8A-C). The first, second, and third distal tarsals (Dt1–3) are visible, although the boundary between the first and second distal tarsal is difficult to identify precisely. The matrix filling the space between the tarsals (and all bones in the pes) is of a very similar colour and texture to the bones themselves, making identifying the precise boundaries of each bone difficult without further preparation of the fossil.

**Fig. 8.**
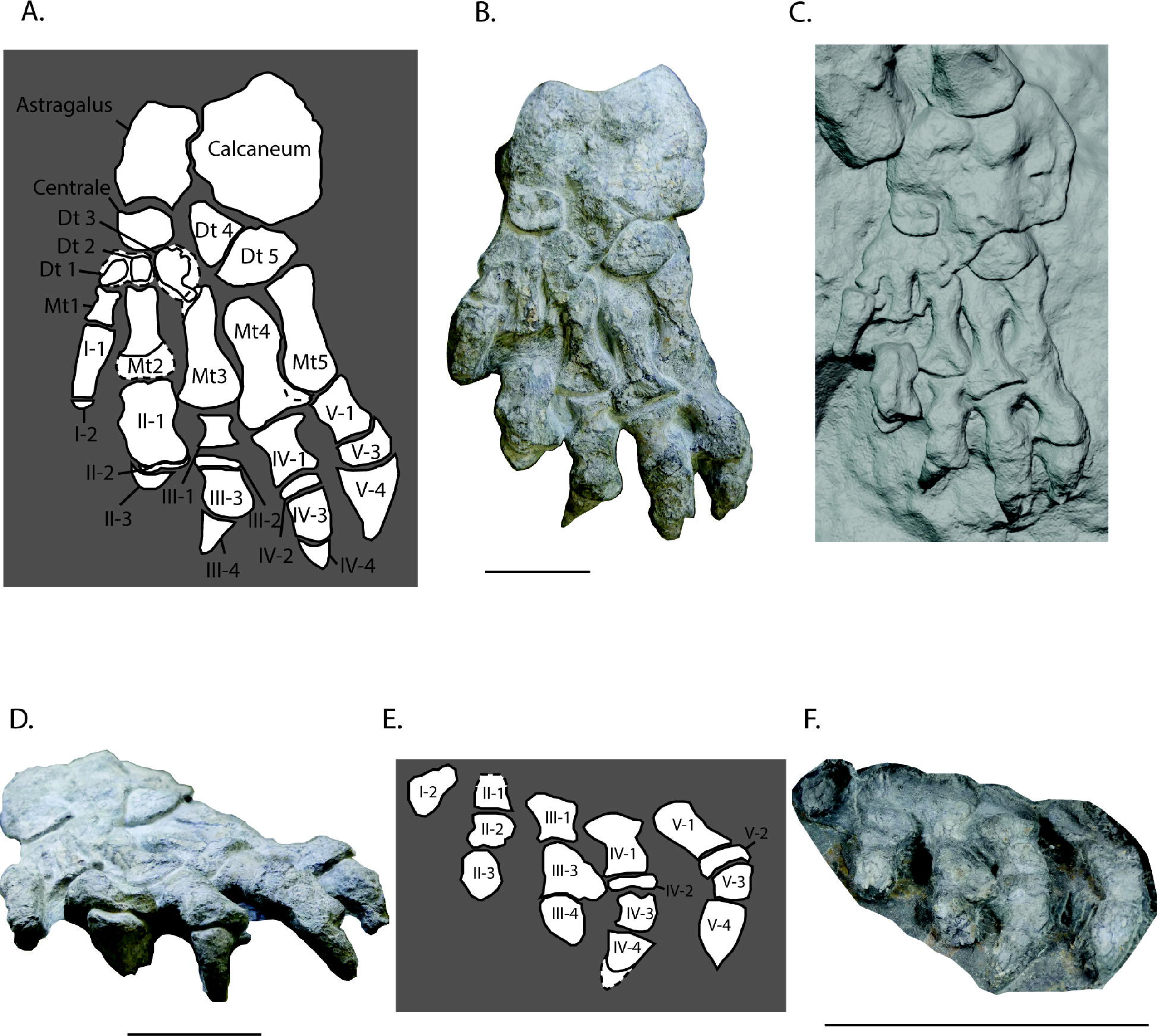
Labelled line drawings (A, E), photographs (B, D, F), and a rendering of a photogrammetric model (C) of the left (A-D) and right (E-F) pes of *Aelurognathus tigriceps* (SAM-PK-K10496) in dorsal (A-C, E-F) and distal (D) views. Dt 4 and Dt 5 comprise the cuboid. The dashed lines infilled with white space represent areas of either bone erosion or bone overlapping bone and the hypothesized full extent of the bone as would have been seen in life. Scale bars for the left pes are both three centimetres, and the scale bar for the right pes is eight centimetres. Abbreviations: Dt = Distal tarsal, Mt = Metatarsal.

The astragalus is rectangular with a convexly curved lateral margin, and overlaps the calcaneum proximally (Fig. 8B) as has been described for other gorgonopsians including *Lycaenops ornatus* (AMNH FARB 2240) and *Gorgonops torvus* (SAM-PK-K10591), albeit to a lesser extent in SAM-PK-K10496 (Colbert 1948; Bendel et al. 2023). The astragalus articulates with the both the fibula and tibia (Fig. 6B) via proximolaterally- and proximomedially-oriented facets at approximately 90° to each other. The facet for the tibia is larger and occupies much of the medial surface of the astragalus. The distal surface of the astragalus forms a broad and concave contact with the centrale, with a small distolateral facet for Dt4 (Fig. 8B). The mediolateral width of the astragalus is greater proximally than distally, similar to *Gorgonops torvus* (SAM-PK-K10591) and *Lycaenops ornatus* (AMNH FARB 2240; in which the difference is larger) (Colbert 1948; Bendel et al. 2023). This differs from the equal proximal and distal widths seen in *Gorgonops* sp. (BSPG 1934 VIII 28) and indeterminate gorgonopsian NHCC LB1073,) (Broili and Schröder 1935; Colbert 1948; Sidor 2022; Bendel et al. 2023).

The calcaneum is very large relative to the other tarsals, at more than double the size of the astragalus (Table 2), similar to *Lycaenops ornatus* (AMNH FARB 2240) and the inferred size of indeterminate gorgonopsian NHCC LB1073, but relatively larger than in *Gorgonops torvus* (SAM-PK-K10591) (Colbert 1948; Sidor 2022; Bendel et al. 2023). The calcaneum is generally hexagonal, with a slightly concave, proximomedially-facing articular surface for the fibula (Fig. 8B). The location and concavity of the fibular articular surface is very similar to that of *Arctops umulunshi* (NHCC LB369) (Mann and Sidor 2025). The medial surface, which articulates with the astragalus, is also slightly convex. Distally, the calcaneum contacts the cuboid. The calcaneum is mediolaterally wider than in *Gorgonops torvus* (SAM-PK-K10591) and *Arctops umulunshi* (NHCC LB369), more closely resembling the width of *Dinogorgon rubidgei* (BP/1/2167) and *Lycaenops ornatus* (AMNH FARB 2240) (Colbert 1948; Sigogneau 1970; Bendel et al. 2023; Mann and Sidor 2025). The exposed surface of the calcaneum is weakly rugose, particularly around its contact with the fourth distal tarsal.

**Table 2.** Lengths and widths of the tarsals in the left pes of *Aelurognathus tigriceps* (SAM-PK-K10496). All measurements are in millimetres.

| Element | Maximum proximodistal length | Maximum mediolateral width |
| --- | --- | --- |
| Calcaneum | 44.5 | 37.1 |
| Astragalus | 28.8 | 17.4 |
| Centrale | 11.2 | 16.2 |
| Distal tarsal 1 | 8.5 | 14.3 |
| Distal tarsal 2 | 16.3 | 9.7 |
| Distal tarsal 3 | 17.8 | 10.9 |
| Cuboid (Distal tarsals 4+5) | 17.0 | 28.6 |

The centrale is rectangular with rounded corners, being slightly wider mediolaterally than long proximodistally (Table 2). It forms a broad, convex contact with the astragalus proximally, a straight contact with Dt4 laterally, and approximately equal-sized contacts with Dt1–3 distally (Fig. 8B). The relative size of the centrale (proximodistal length) is similar to those of *Dinogorgon rubidgei* (BP/1/2167) and *Lycaenops ornatus* (AMNH FARB 2240). It is proportionally larger than those of some other gorgonopsians, such as *Gorgonops* sp. (BSPG 1934 VIII 28), *Gorgonops torvus* (SAM-PK-K10591) and indeterminate gorgonopsian NHCC LB1073 (Broili and Schröder 1935; Colbert 1948; Sigogneau 1970; Sidor 2022; Bendel et al. 2023).

The first and second distal tarsals are small, around two-thirds the diameters of the centrale (Fig. 8B; Table 2), similar to that of *Gorgonops torvus* (SAM-PK-K10591), but smaller than those of *Dinogorgon rubidgei* (BP/1/2167), *Lycaenops ornatus* (AMNH FARB 2240), and indeterminate gorgonopsian NHCC LB1073 (Broili and Schröder 1935; Colbert 1948; Sigogneau 1970; Sidor 2022; Bendel et al. 2023). Dt1 is oval with a proximolateral-to-distomedial oriented long axis and contacts the first metatarsal (Mt1) distally (Fig. 8B). This is the same orientation seen in that of *Gorgonops torvus* (SAM-PK-K10591) whereas in *Gorgonops* sp. (BSPG 1934 VIII 28) the long axis of Dt1 is oriented medial-to-laterally. Dt2 is square and contacts the second metatarsal (Mt2) distally and the centrale proximally (Fig. 8B). The small, blocky shape of Dt2 closely resembles those of indeterminate gorgonopsian NHCC LB1073 and *Gorgonops torvus* (SAM-PK-10591).

Dt3 is larger than Dt1 and Dt2, and is similar in size to the centrale (Table 2), unlike in indeterminate gorgonopsian NHCC LB1073 where Dt1 is the largest of the first three distal tarsals (Sidor 2022); however, this apparent difference in size may be a result of incomplete preparation, or erosion of Dt1. The larger size of Dt3 more closely resembles that of *Gorgonops* sp. (BSPG 1934 VIII 28), *Lycaenops ornatus* (AMH FARB 2240), and *Dinogorgon rubidgei* (BP/1/2167) (Broili and Schröder 1935; Colbert 1948; Sigogneau 1970). The overall shape of the third distal tarsal is irregular; it has a curved proximal portion, from which a slender distolateral process emerges to contact the proximal end of Mt3 (Fig. 8B). However, this unusual shape is likely the result of damage, as it does not resemble the elongate, approximately parallelogram shape seen in numerous other gorgonopsians such as *Lycaenops ornatus* (AMNH FARB 2240), *Gorgonops* sp. (BSPG 1934 VIII 28), and *Dinogorgon rubidgei* (BP/1/2167), nor does it resemble the teardrop shape seen in *Gorgonops torvus* (SAM-PK-K10591) and indeterminate gorgonopsian NHCC LB1073 (Broili and Schröder 1935; Colbert 1948; Sigogneau 1970; Sidor 2022; Bendel et al. 2023). Dt3 contacts the centrale proximally, the second distal tarsal medially, the cuboid laterally, and the third metatarsal distally.

The cuboid (combined Dt4–5) has a semilunate outline (Fig. 8B), as in most other known gorgonopsians, such as those of *Lycaenops ornatus* (AMNH FARB 2240), *Dinogorgon rubidgei* (BP/1/2167), *Gorgonops torvus* (SAM-PK-K10591) and indeterminate gorgonopsian NHCC LB1073, with the concave edge facing proximolaterally and contacting the calcaneum (Colbert 1948; Sigogneau 1970; Sidor 2022; Bendel et al. 2023). This differs only from the condition seen in *Gorgonops* sp. (BSPG 1934 VIII 28) where the cuboid is more triangular with concave medial and distal edges and a convex edge contacting the calcaneum (Broili and Schröder 1935). The dorsal surface of the cuboid is mostly flat, but the lateral third of Dt5 slopes ventrolaterally. The cuboid makes a small proximomedial contact with the astragalus, and contacts the centrale medially, Dt3 distomedially, and Mt 4–5 distally. The contact between Dt5 and Mt5 is extensive, covering most of the proximal end of Mt5 (Fig. 8B). This extent of contact is similar to that of *Gorgonops torvus* (SAM-PK-K10591), *Dinogorgon rubidgei* (BP/1/2167) and *Lycaenops ornatus* (AMNH FARB 2240), as opposed to the more minimal contact seen in *Gorgonops* sp. (BSPG 1934 VIII 28) and indeterminate gorgonopsian NHCC LB1073. That there is a difference in this contact between two specimens of the same genus (SAM-PK-K10591 and BSPG 1934 VIII 28) suggests a greater potential for individual variation over phylogenetic patterns here. A prominent fissure through the cuboid separates it into two subequal portions presumably corresponding to Dt4 and Dt5 (Fig. 8B). This fissure is absent in all other known gorgonopsian cuboids, including *Lycaenops ornatus* (AMNH FARB 2240), *Gorgonops* sp. (BSPG 1934 VIII 28), *Dinogorgon rubidgei* (BP/1/2167), *Gorgonops torvus* (SAM-PK-K10591), and indeterminate gorgonopsian NHCC LB1073, and as all these specimens have been interpreted as adults, its presence in SAM-PK-K10496 suggests that fusion of these elements occurred relatively late in ontogeny in *Aelurognathus* (Broili and Schröder 1935; Colbert 1948; Sigogneau 1970; Sidor 2022; Bendel et al. 2023).

### Metatarsals

All five left metatarsals are preserved and exposed (Fig. 8A-C), whereas the right metatarsals are likely preserved, but are not exposed. The metatarsals increase in proximodistal length from medial to lateral, with the Mt1 being the shortest (10.0 mm) by far (compared to 23.9 mm in Mt2, including an estimated 5.8 mm of missing Mt2 preserved as impression) and Mt5 (36.4 mm) being the longest (Table 3). All the metatarsals are constricted around midlength (Table 3). Mt1 is about as proximodistally long as it is mediolaterally wide, with a minimum shaft width of 3.7 mm, forming a barbell-shaped outline in anterior view (Fig. 8B). The proximal and distal ends of Mt1 are concave, forming articular facets for the first distal tarsal proximally and the first proximal phalanx distally (Fig. 8B).

**Table 3.**
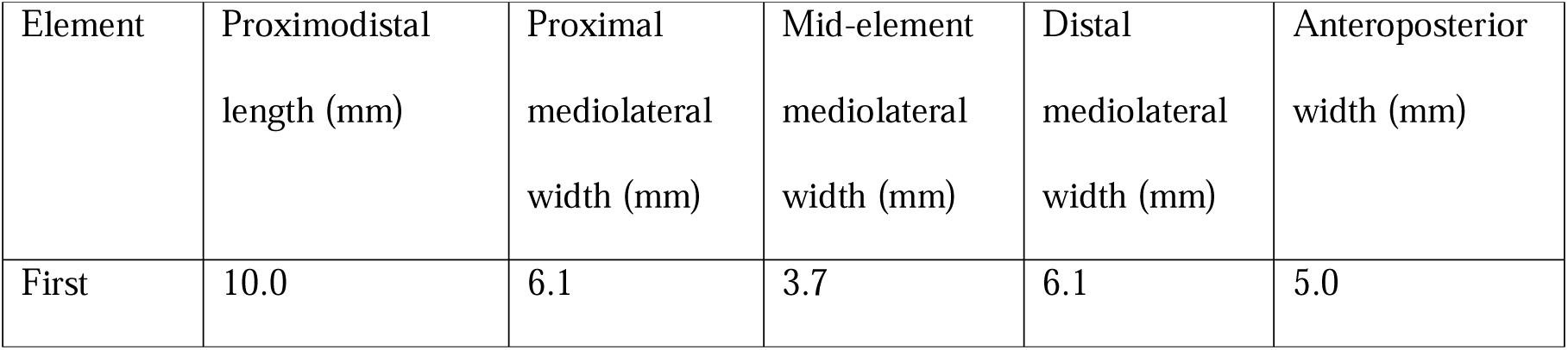
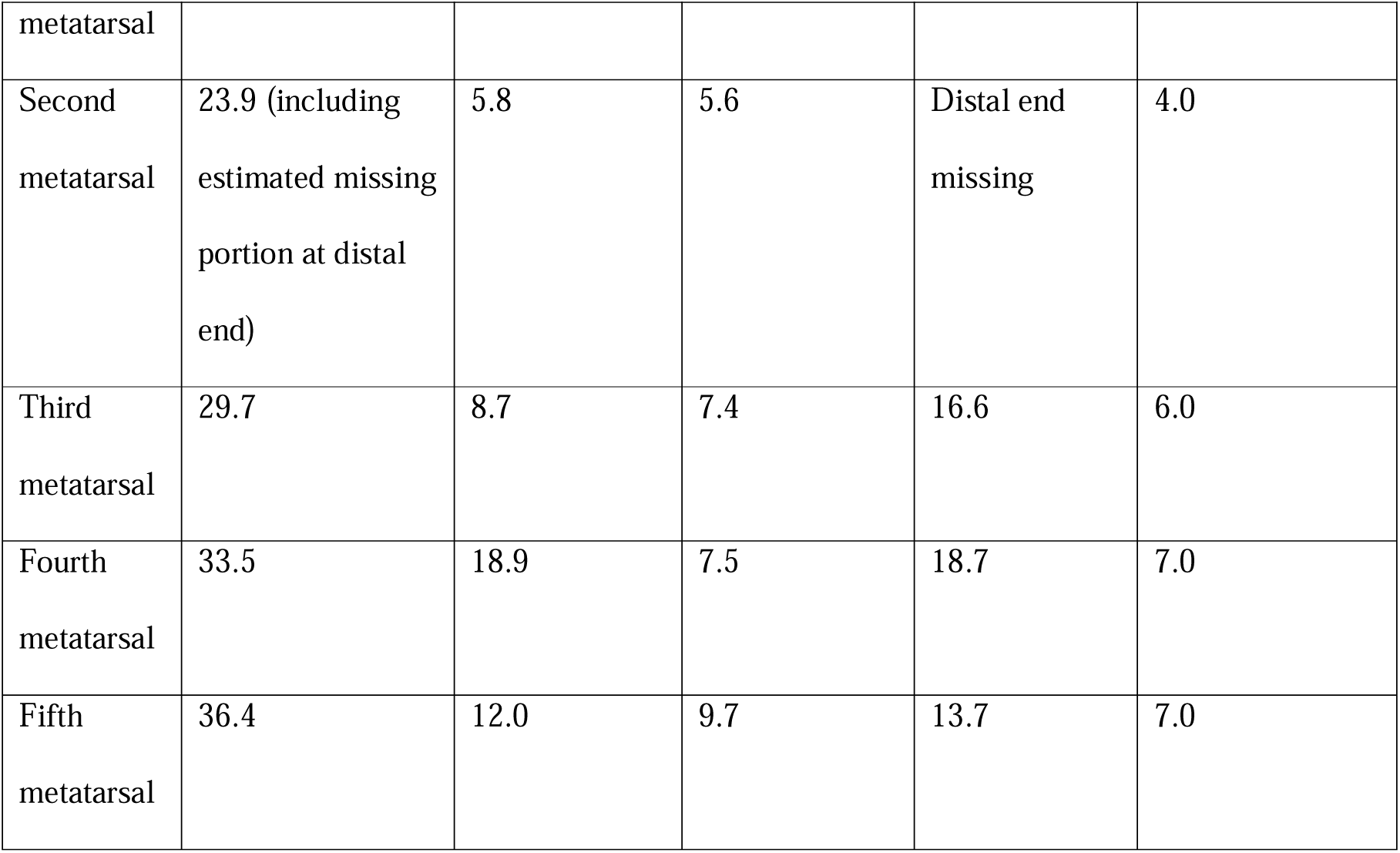
Length and width measurements of the metatarsals in the left pes of *Aelurognathus tigriceps* (SAM-PK-K10496). All measurements are in millimetres.

The distal end of Mt2 is slightly wider than the proximal end (Table 3). It articulates with Dt1 and Dt2 at the proximal end, and would articulate with the second proximal phalanx (II-1) at the distal end if it was completely preserved.

Mt3 is large with a proximodistal length of 29.7 mm (Table 3). The overall shape of Mt3 is proximally narrow with a wider, flared distal end (Fig. 8B). A fossa is present on the dorsal distal surface of Mt3 (Fig. 8C), as also seen in *Gorgonops torvus* (SAM-PK-K10591), but is absent from other specimens such as *Gorgonops* sp. (BSPG 1934 VIII 28) and indeterminate gorgonopsian NHCC LB1073 (Broili and Schröder 1935; Sidor 2022; Bendel et al. 2023). Mt3 articulates proximally with the Dt2 and Dt3, and articulates distally with the third proximal phalanx (III-1).

Mt4 is slightly larger than Mt3, with a proximodistal length of 33.5 mm (Fig. 8B). The narrowing along the shaft in Mt4 is more extreme than in Mt3, narrowing by over a factor of two (Table 3). Mt4 articulates with the cuboid proximally and with the fourth proximal phalanx (IV-1) distally (Fig. 8B).

Mt5 is larger and stockier than the other metatarsals, with a proximodistal length of 36.4 mm and a mediolateral width of 9.7 mm at the midshaft (Table 3). The overall shape of Mt5 also has a barbell shape, being wider at both the proximal and distal ends, and narrower in the middle (Fig. 8B). The midshaft narrowing on Mt5 is by about one-third (Table 3). Mt5 articulates with the cuboid proximally and the fifth proximal phalanx (V-1) distally (Fig. 8B).

### Pedal phalanges

All phalanges on the left foot except the fifth disc phalanx (V-2) are preserved and exposed (Fig. 8A-D), and on the right foot all phalanges except the first proximal phalanx (I-1) and the third disc phalanx (III-2) are preserved and exposed (Fig. 8E-F). The pedal phalangeal formula is 2-3-4-4-4, based on information from both pedes (Fig. 8). This differs from the pedal phalangeal formula of other specimens such as indeterminate gorgonopsian NHCC LB1073 (2-3-4-5-4), and the *Gorgonops torvus* specimen SAM-PK-K10591 (2-3-4-5-3) (Sidor 2022; Bendel et al. 2023). In the left foot, the phalanges are preserved with the dorsal side exposed (Fig. 8A-D), whereas in the right foot the phalanges are preserved with the ventral side exposed (Fig. 8E-F). In the left foot, all ungual phalanges are flexed to an angle of 90° relative to the non-ungual phalanges, except the fifth ungual phalanx (V-4) (Fig. 8D).

Other than the disc phalanges (III-2, IV-2, V-2), non-ungual phalanges (I-1, II-1, II-2, III-1, III-3, IV-1, IV-3, V-1, V-3) share a similar morphology to that of the metatarsals, with a barbell-like shape that is proximally and distally wider than at the mid-element (Fig. 8B). Other than I-1 (first proximal phalanx), the phalanges are proximodistally shorter than their metatarsals (Tables 3–4). The second proximal phalanx (II-1) is the largest, being mediolaterally wider than its corresponding metatarsal (Table 4).

**Table 4.**
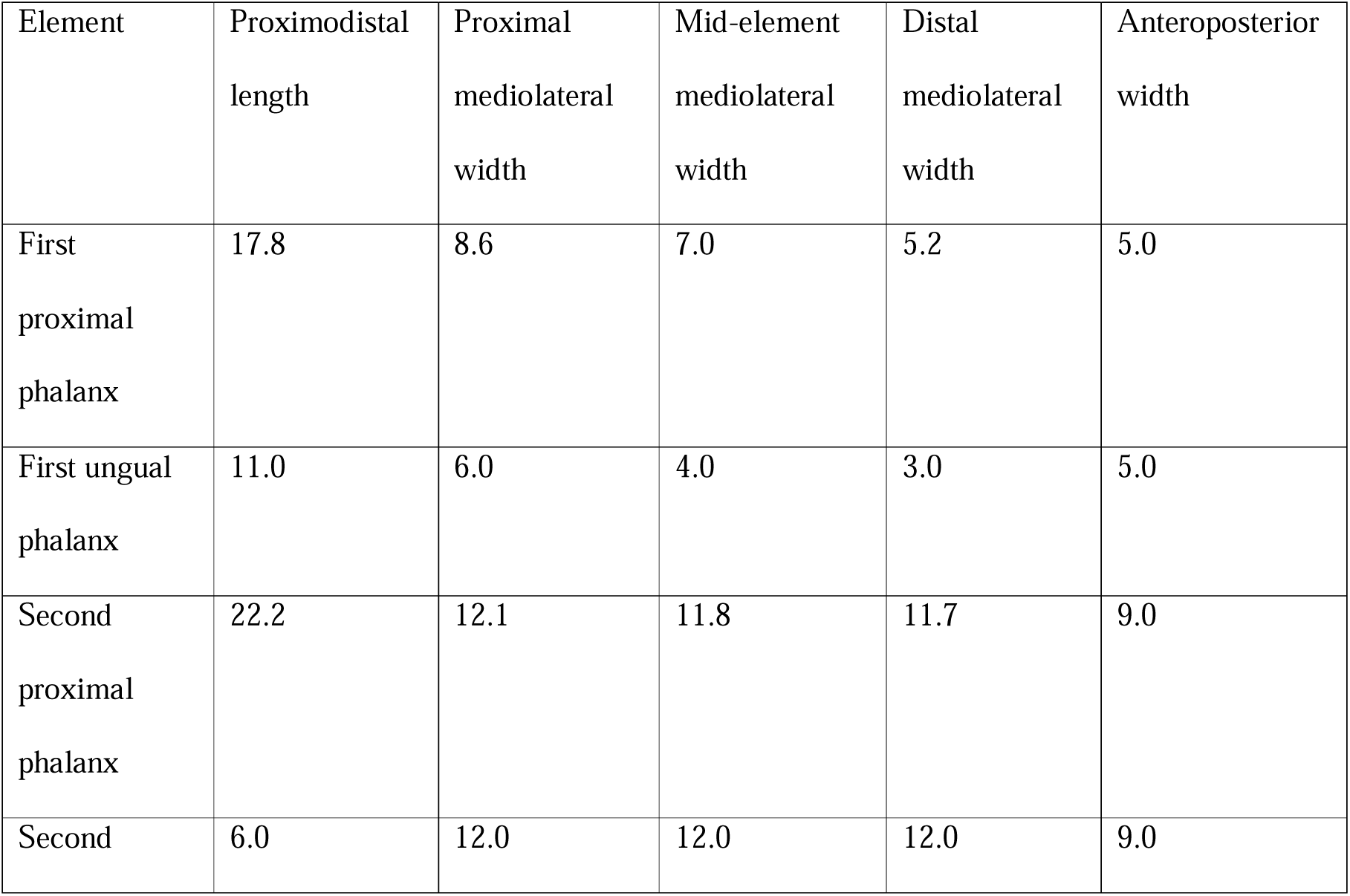
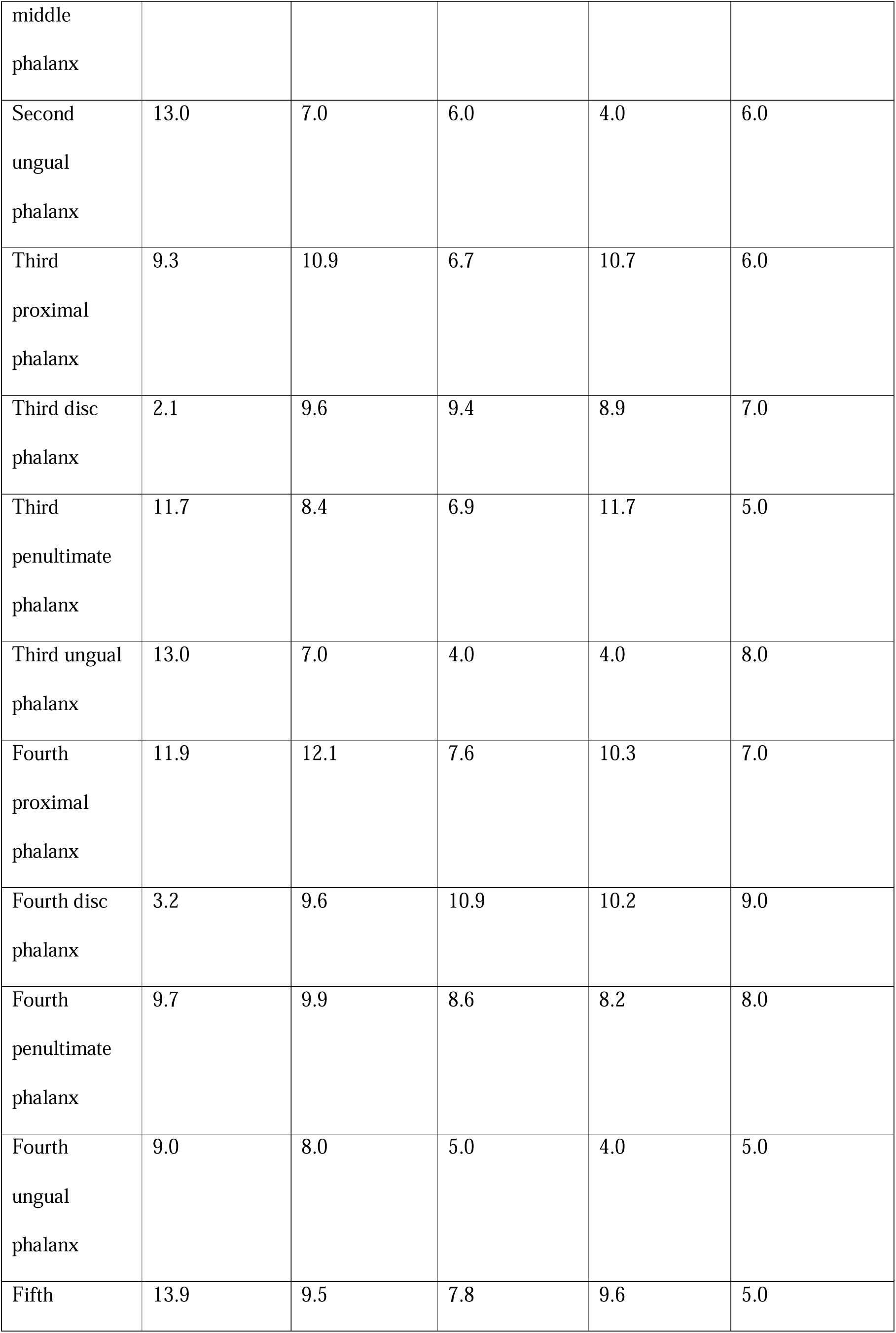
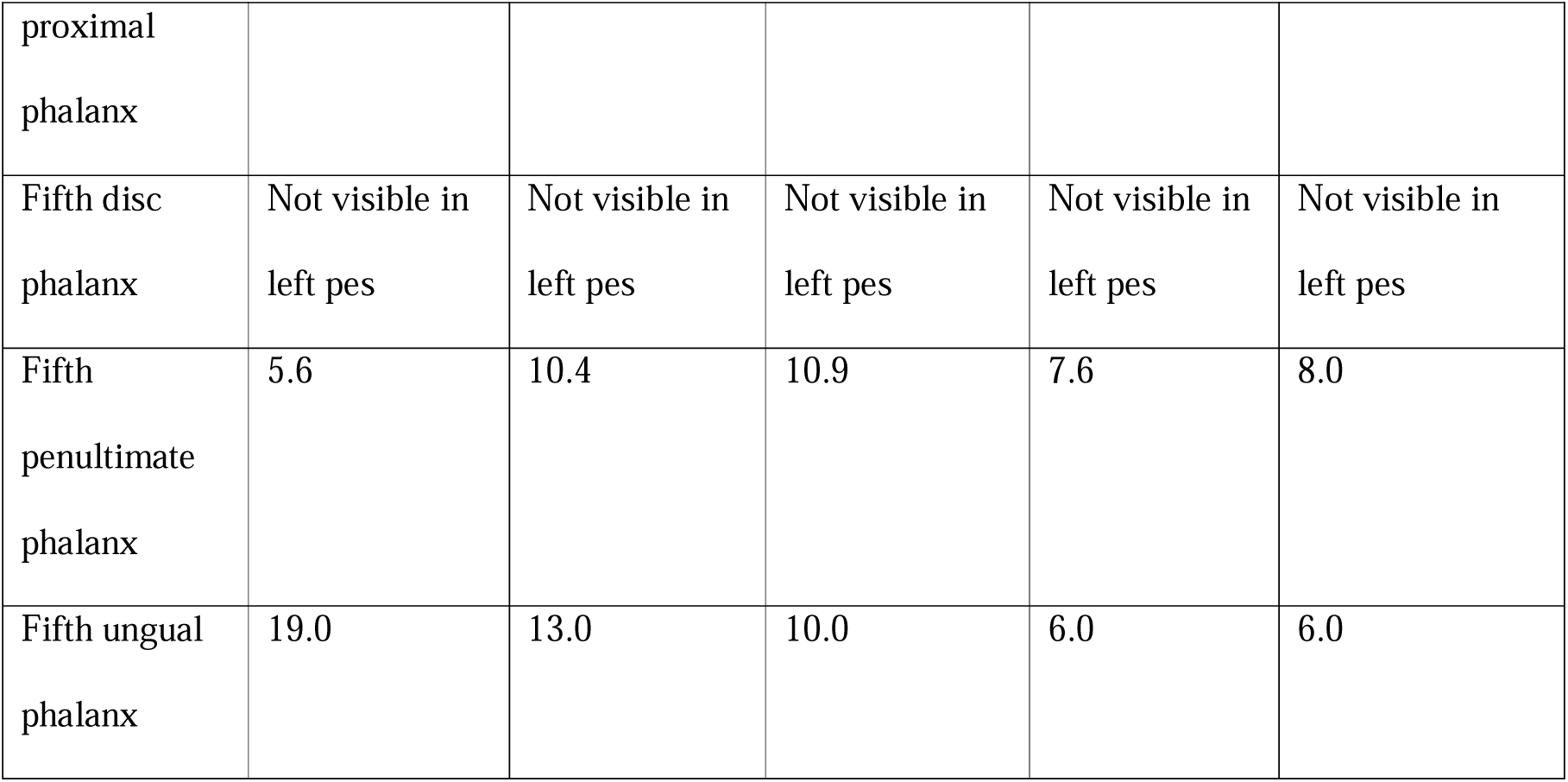
Length and width measurements for pedal phalanges in the left pes of *Aelurognathus tigriceps* (SAM-PK-K10496). All measurements are in millimetres.

Disc phalanges are present in the third–fifth digits (III-2, IV-2, V-2), and are proximodistally short and mediolaterally wide (Table 4). The reduced pedal phalangeal formula of SAM-PK-K10496 results from the absence of one disc phalanx from digit 4, (compared to e.g. indeterminate gorgonopsian NHCC LB1073, *Gorgonops torvus* (SAM-PK-10591), and *Dinogorgon rubidgei* (BP/1/2167); (Sigogneau-Russell 1989; Sidor 2022; Bendel et al. 2023)). This phalanx appears to be absent in both the left and right pes, and its absence in the right pes is more clearly evidenced because the space between adjacent phalanges, where it would otherwise be present, is more exposed (Fig. 8B, E).

The pedal ungual phalanges are triangular, and curve ventrodistally, tapering to a point terminally (Fig. 8B, D-E). The ungual phalanges are semilunar in cross-section, with a convex dorsal surface and concave ventral surface.

## Discussion

The pectoral girdle of SAM-PK-K10496 is the most fragmentary part of the skeleton, making a clear interpretation difficult. Fortunately, the holotype of *Aelurognathus tigriceps* (SAM-PK-2342) includes a complete left pectoral girdle that can be used as a point of comparison. The pectoral girdle in SAM-PK-2342 shows a large, curved scapulocoracoid, with a smaller, shallow interclavicle positioned medially, and, anteriorly, a clavicle connecting the anterior edge of the interclavicle to the anteriormost point of the scapula (Broom and Haughton 1913). In SAM-PK-10496, the most immediately comparable element is the partially preserved scapulocoracoid, as a result of its distinct curved shape and still remaining in articulation with the rest of the skeleton. However, the other pectoral elements cannot be identified.

Variation in the pedal phalangeal formula of gorgonopsians is poorly-documented due to the scarcity of complete or near-complete gorgonopsian pedes that have been described (Sigogneau 1970; Sigogneau-Russell 1989; Hopson 1995; Sidor 2022; Bendel et al. 2023). Prior inferences of pedal phalangeal formula in gorgonopsians has largely come from either comparison to the manus (Rowe and Van Den Heever 1986; Hopson 1995) or from a handful of more recent complete specimens that have been used to make the best possible broad inferences given the limited information available (Sidor 2022; Bendel et al. 2023). The manual phalangeal formula is 2-3-4-5-3 in most gorgonopsians reported thus far (Hopson 1995; Kümmell and Frey 2014; Bendel et al. 2023), which fits within the hypothesized range of pedal phalangeal formulae of *Viatkogorgon ivakhnenkoi* (PIN 2212/61) (Tatarinov 2004; Bendel et al. 2023). However, the indeterminate gorgonopsian NHCC LB1073 has a clear phalangeal formula of 2-3-4-5-4 (Sidor 2022), matching the known digits for *Arctops umulunshi* (NHCC LB396) with a formula of ?-3-4-5-4 (Mann and Sidor 2025), and indicating some variability of pedal phalangeal formula among gorgonopsians. SAM-PK-K10496 further adds to this variability with a phalangeal formula of 2-3-4-4-4, with only one disc-like phalanx in the fourth digit, as opposed to the two seen in indeterminate gorgonopsian NHCC LB1073, *Gorgonops torvus* (SAM-PK-K10591), or *Arctops umulunshi* (NHCC LB396) (Sidor 2022; Bendel et al. 2023; Mann and Sidor 2025). Bendel et al. (2023) used the differences between the phalangeal formula of NHCC LB1073 and *Gorgonops torvus* SAM-PK-K10591 to suggest that there may have been reductions in more later groups, including rubidgeines. As a rubidgeine, the reduced phalangeal count of the fourth digit of SAM-PK-K10496 is consistent with this prediction.

SAM-PK-K10496 potentially represents a subadult, based on incomplete fusion of the cuboid (Dt4+Dt5) and smaller size (skull height at orbit = 10.4 cm; preserved skull length = 13.8 cm) compared to the holotype of *Aelurognathus tigriceps* (SAM-PK-2342; skull height = 18.8 cm; and 22.2 cm for the equivalent length measurement, from the tip of the snout to the ventral orbital mid-length).

SAM-PK-K10496 includes one of the most complete and best-preserved gorgonopsian tails reported thus far, and is the longest gorgonopsian tail currently described in terms of number of caudal vertebrae present. Other relatively complete tails have been reported in *Viatkogorgon ivakhnenkoi* (PIN 2212/61) and unidentified Tanzanian gorgonopsian UMZC T883. However, detailed descriptions in both of these cases are lacking – prior literature has referenced the need for a more detailed postcranial description of PIN 2212/61 to complement its initial preliminary description (Tatarinov 2004; Kammerer and Masyutin 2018), and UMZC T883 has yet to be described, despite having been referenced in comparison to other specimens numerous times (Kemp 1969; Sigogneau-Russell 1989; Bishop et al. 2022; Bendel et al. 2023; Mann and Sidor 2025).

SAM-PK-K10496 has at least 23 caudal vertebrae, whereas *Viatkogorgon ivakhnenkoi* (PIN 2212/61) and indeterminate gorgonopsian UMZC T883 have a maximum caudal vertebral count of 20 (Tatarinov 2004; SKP pers. obs). Outside of gorgonopsians, caudal vertebral count in non-mammalian therapsids varies from group to group. Among the Biarmosuchia, extensive caudal material is known only from *Hipposaurus boonstrai* (SAM-PK-8950, SAM-PK-12010), with an estimated caudal count of at least 26 (Sigogneau-Russell 1989) which suggests a similar, if somewhat longer, tail length than that of SAM-PK-K10496. Among the Dinocephalia, only two fairly complete tails are known: the anteosaur *Titanophoneus potens* (PIN 157/1), which is noted to have at least 38 caudal vertebrae but likely up to 60 (Orlov 1958), and the tapinocephalian *Tapinocaninus pamelae* (NMQR 2987), which has a total of seven caudal vertebrae (Rubidge et al. 2019), indicating extreme disparity in tail length between subclades in this group. Anomodontia also shows differing tail lengths between different subgroups – basal anomodonts are noted as having long tails, such as in the venyukovioid *Suminia getmanovi* (PIN 2212/116) with at least 52 caudal vertebrae (Fröbisch and Reisz 2011), whereas derived dicynodonts have much shorter tails that range between eight and 14 vertebrae (Ray and Chinsamy 2003; Maharaj et al. 2021; Xie et al. 2023). Therocephalians are generally noted as having very short tails, such as in UMZC T837, which has maximally 10 caudal vertebrae (Kemp 1986; Huttenlocker et al. 2022). When compared to other therapsids, SAM-PK-K10496 represents an intermediate tail length, between the extreme highs of the basal anomodonts and anteosaurs and the extreme lows of the dicynodonts and therocephalians. While prior literature does not often discuss caudal material in therapsids, sometimes referring to a therapsid condition “very much reduced in length compared to pelycosaurs” as in Kemp (2005), there as yet has not been a rigorous comparison of tail lengths across Therapsida, with only a handful of published specimens containing notable caudal material. However, the presence of a longer tail in SAM-PK-K10496 raises the possibility of a primitively longer tail in therapsids, followed by multiple reductions among subgroups including Tapinocephalidae (minimum caudal count = 7 (Rubidge et al. 2019)), Dicynodontia (minimum caudal count = 8 (Ray and Chinsamy 2003; Maharaj et al. 2021; Xie et al. 2023)), and Therocephalia (minimum caudal count = 9 (Kemp 1986, 2005)).

## Conclusion

1. SAM-PK-K10496 is a subadult specimen of *Aelurognathus tigriceps* that preserves new data for the posterior half of the skeleton.
2. The complete pes gives a pedal phalangeal formula of 2-3-4-4-4 which adds to the variability in pedal phalangeal formula known for gorgonopsians.
3. At between 23 and 27 caudal vertebrae, the tail contains more segments than any other known gorgonopsian, and may have represented an intermediate or primitively longer condition among therapsids more broadly.

## Acknowledgements

We would like to thank Zaituna Skosan for providing access to the Iziko South African Museums collections, and Roger M. H. Smith for providing detailed locality information and field notes associated with the specimen. We are also grateful to Matt Lowe (UMZC) and Bernhard Zipfel (BP) for providing access to comparative materials. SKP was funded by NERC Award NE/S007474/1. The funders had no role in study design, data collection and analysis, decision to publish, or preparation of the manuscript. We would also like to thank Christian Sidor and Adam Huttenlocker for their helpful reviews and comments which have significantly improved this work.

